# Lignin-based resistance to *Cuscuta campestris* parasitism in Heinz resistant tomato cultivars

**DOI:** 10.1101/706861

**Authors:** Min-Yao Jhu, Moran Farhi, Li Wang, Richard N. Philbrook, Michael S. Belcher, Hokuto Nakayama, Kristina S. Zumstein, Steven D. Rowland, Mily Ron, Patrick M. Shih, Neelima R. Sinha

## Abstract

*Cuscuta* species (dodders) are agriculturally destructive parasitic angiosperms. These parasitic plants use haustoria as physiological bridges to extract nutrients and water from hosts. *Cuscuta campestris* has a broad host range and wide geographical distribution. While some wild tomato relatives are resistant, cultivated tomatoes are generally susceptible to *C. campestris* infestations. However, some specific Heinz tomato hybrid cultivars exhibit resistance to dodders in the field, but their defense mechanism was unknown. Here, we discovered that the stem cortex in these resistant lines responds with local lignification upon *C. campestris* attachment, preventing parasite entry into the host. *LIF1* (*Lignin Induction Factor 1*, an *AP2*-like transcription factor), *SlMYB55*, and *CuRLR1* (*Cuscuta R-gene for Lignin-based Resistance 1*, a *CC-NBS-LRR*) are identified as crucial factors conferring host resistance by regulating lignification. *SlWRKY16* is upregulated upon *C. campestris* infestation and acts as a negative regulator of *LIF1* function. Intriguingly, *CuRLR1* may play a role in signaling or function as a receptor for receiving *Cuscuta* signals or effectors to regulate lignification-based resistance. In summary, these four regulators control the lignin-based resistance response, preventing *C. campestris* from parasitizing these resistant tomatoes. This discovery provides a foundation for investigating multilayer resistance against *Cuscuta* species and has potential for application in other essential crops attacked by parasitic plants.

**One-sentence summary:** Four key regulators confer lignin accumulation in the tomato stem cortex to block *C. campestris* host penetration upon infection.

## Introduction

Parasitic plants directly attach to hosts using specialized haustorial organs known as haustoria. These connections function as physiological bridges to extract nutrients and water from the hosts, making most traditional herbicides and control methods, including management of soil fertility, hand weeding, and sanitation, either too cost-intensive, labor-intensive, or ineffective in regulating parasitic plant infestations. Therefore, parasitic angiosperms are among the most devastating pests, reducing the yields of agricultural crops each year by billions of dollars worldwide (Agrios, 2005; Yoder and Scholes, 2010). Members of the *Cuscuta* genus (family Convolvulaceae), also known as dodders, occur worldwide and *Cuscuta* infestations in tomato alone lead to 50–72% yield reductions (Yaakov et al., 2001). Despite serious agricultural problems caused by *Cuscuta*, our understanding of the interactions between *Cuscuta* and its hosts is relatively limited compared to our knowledge of pathogenic fungi, bacteria, and viruses. Only recently, the first receptor (*CuRe1*, Solyc08g016270), an LRR receptor-like serine/threonine-protein kinase (RLP), from *Cuscuta* was identified in tomatoes (Hegenauer et al., 2016; Hegenauer et al., 2020). *CuRe1* initiates PAMP (Pathogen-associated molecular pattern)-triggered immunity (PTI) to *Cuscuta reflexa*. However, plants that lack *CuRe1* are still fully resistant to *C. reflexa*. This result indicates that other layers of defense, besides *CuRe1,* must also be involved in the responses to these parasites. Therefore, further investigating the potential multilayered resistance mechanisms will aid in developing parasitic plant-resistant crops.

Potential immune responses and defense responses to parasitic plants have been observed in several crop plants, including rice, tomato, cowpea, and sunflower (Mutuku et al., 2015; Hegenauer et al., 2016; Duriez et al., 2019; Su et al., 2020). Notably, most previous reports indicated that hypersensitive response is the major mechanism that contributes to plant immunity to parasitic plants (LANE et al., 1993; Hegenauer et al., 2016; Su et al., 2020). A few studies indicated that secondary cell-wall modification and formation also contribute to defense in root parasitic plants (Yoshida and Shirasu, 2009; Mutuku et al., 2019). Plants often modify their cell walls in response to pathogen infection and herbivore feeding (Moura et al., 2010). Among different modifications, lignification is considered a major mechanism for resistance in plants (Vance et al., 1980; Moura et al., 2010; Bellincampi et al., 2014; Malinovsky et al., 2014). Lignified cell walls have higher mechanical strength, are impermeable to water, and less accessible to cell wall-degrading enzymes (Bhuiyan et al., 2009; Barros et al., 2015). Several previous reports indicated that lignified endodermal cells were found in resistant host roots, like vetch (*Vicia* spp.) and faba bean (*Vicia faba*), in response to root parasitic plant attack (PÉRez-De-Luque et al., 2005; Pérez-de-Luque et al., 2007). However, how secondary cell-wall modification and lignin are involved in the defense responses to stem parasitic plants still needs to be elucidated. Thus, we specifically investigated host cell wall composition and the lignin biosynthesis pathways aiming to discover the potential additional layers of resistance to *Cuscuta* spp.

*Cuscuta campestris* (*C. campestris*) attacks a wide range of crop species worldwide (Lanini and Kogan, 2005). Although cultivated tomatoes are usually susceptible (Ashton, 1976), some specific Heinz tomato cultivars that are resistant to *Cuscuta* spp. were discovered in the field (Hembree et al., 1999; Yaakov et al., 2001). These resistant cultivars have been used in the field to control the infestation of *Cuscuta* spp., but the resistance mechanism remains unknown. Therefore, to identify the underlying mechanism and genes involved in these defense responses, these dodder-resistant Heinz tomatoes were used for further study. We discovered that the resistance response in these Heinz cultivars is based on post-attachment lignification in the stem cortex upon *C. campestris* infection. Recent work described the involvement of lignin in the resistance responses to root parasitic plants (Labrousse et al., 2001; Kusumoto et al., 2007; Cui et al., 2018). However, considering the differences in the anatomy of stems and roots, whether host plants deploy similar mechanisms to stop stem parasitic plants remains under-investigated. Based on our comparative transcriptomics, virus-induced gene silencing, and gain-of-function studies in susceptible cultivars, we identified two transcription factors, *SlMYB55* (Solyc10g044680) and *LIF1* (Solyc02g079020, *Lignin induction Factor 1*, an *AP2*-like protein), that regulate the biosynthesis of lignin in the cortex. Moreover, *CuRLR1* (Solyc04g009110, a *CC-NBS-LRR*) may be engaged in signaling or function as a receptor for perceiving *C. campestris* signals or effectors, leading to lignification-based resistance. Overexpression of *CuRLR1* in susceptible tomato only induced strong lignification upon *C. campestris* attachment or *C. campestris* extract injection. To investigate whether these newly discovered lignin-based resistance responses connect with previously identified *CuRe1* mediated resistance responses, we conducted comprehensive RNA-Seq clustering and gene-coexpression analysis. Our gene coexpression networks indicate that *CuRe1* is also connected with *CuRLR1, LIF1, and SlMYB55* in resistant cultivars under *C. campestris* infested condition and also helped us identify another transcription factor, *SlWRKY16* (Solyc07g056280), which has a similar expression pattern as *CuRe1*. CRISPR-mediated mutants of *SlWRKY16* showed lignification in the cortex and were more resistant to *C. campestris*. This result indicates that *SlWRKY16* functions as a negative regulator of the lignin-based resistance. Furthermore, we noticed that the lignin-based resistance responds to a large protein molecule from *C. campestris* extracts, which might represent potential novel *C. campestris* signals or effectors. In summary, we discovered four key regulators control the lignin-based resistance response in the stem cortex upon *C. campestris* infection, and this lignification blocks *C. campestris* strands from parasitizing selected Heinz tomato cultivars.

## Results

### Response to *C. campestris* in the resistant cultivars

While most tomato cultivars can be parasitized by *C. campestris*, the Heinz hybrids 9492 and 9553 (H9492 and H9553) exhibit resistance to dodders (Yaakov et al., 2001). *C. campestris* strands grew well on the susceptible H1706 (genome sequenced) and H9775 (Heinz hybrid 9775 – closely related to the resistant cultivars) (Figure 1A). On the other hand, *C. campestris* strands could not form good attachments with H9492 and H9553, and haustoria detached from the host stem, preventing parasite growth (Figure 1B). Based on the biomass ratio of *C. campestris* and host (relative growth rate), H9492 and H9553 cannot support long-term (>45 days) growth of *C. campestris*, in contrast to H9775 and H1706 (Figure 1C).

**Figure 1.**
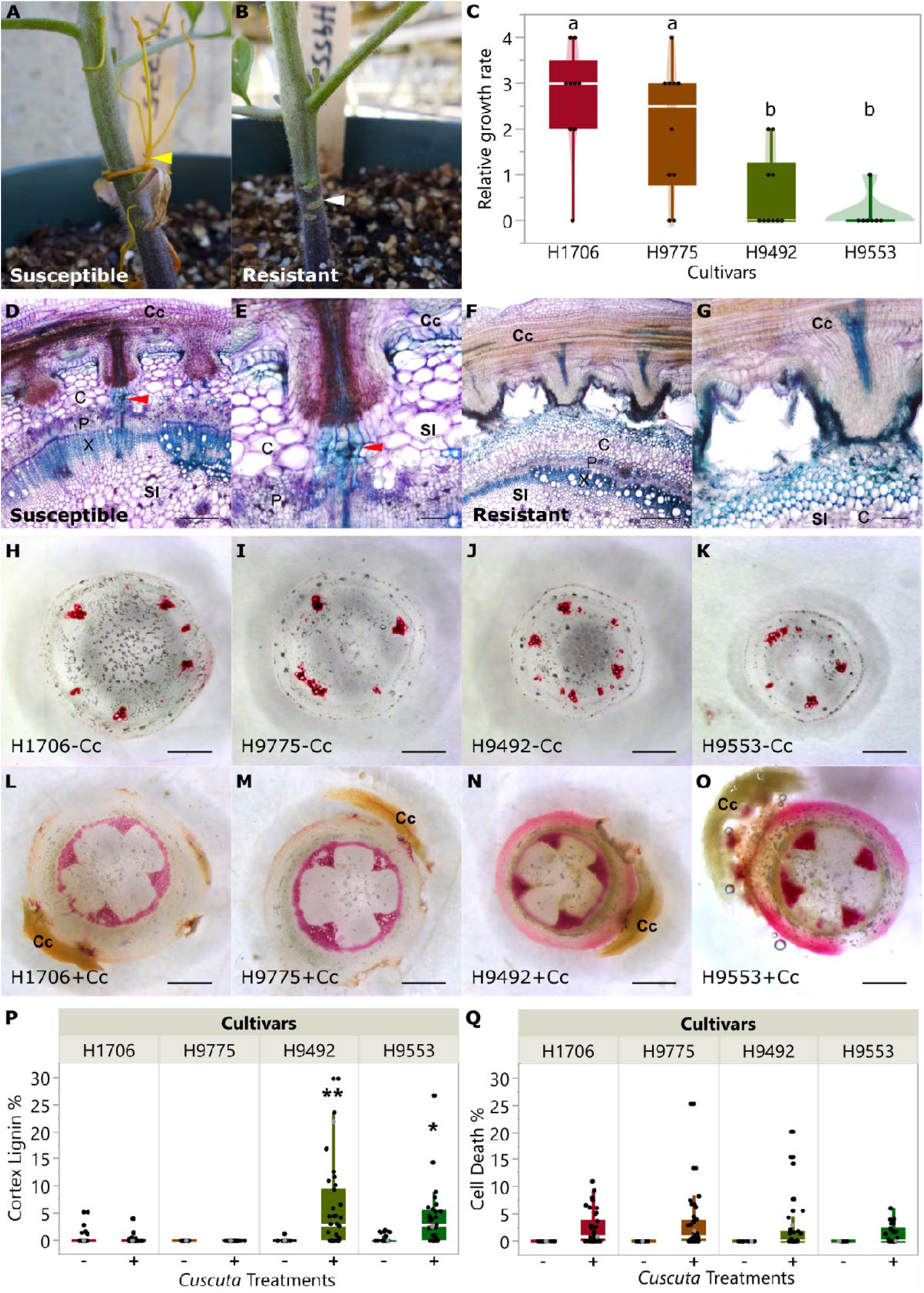
The comparison of resistance responses to *C. campestris* in tomato cultivars. (A) *C. campestris* grows on the susceptible H9775, (B) and cannot attach on the resistant H9553. (C) The biomass ratio of host and *C. campestris* (*Cuscuta* weight/ tomato weight) on different cultivars. Data were assessed using pair-wise comparisons with Tukey test. P-values between “a” and “b” are < 0.05; H1706, n = 9; H9775, n = 10, H9492, n = 10, H9553 n = 7. Data were collected at 45 days post attachment (DPA). (D-G) 100 μm vibratome longitudinal sections of *C. campestris* haustoria attaching to H1706 (D-E) and H9553 (F-G), and stained with Toluidine Blue O. Lignin is stained as blue. Red arrowhead indicates haustorial vascular connections. Cc indicates *C. campestris*; Sl indicates *S. lycopersicum.* C, cortex; P, phloem; X, xylem. (D and F) Scale bar, 40 µm. (E and G) Scale bar, 10 µm. (H-O) are ∼300 μm sections of the haustoria attachment sites stained with Phloroglucinol-HCl. Scale bar, 1 mm. Lignin is stained as red. Stem cross-sections of H1706 (H and L), H9775 (I and M), H9492 (J and N), and H9553 (K and O) without *C. campestris* treatment (labeled with -Cc) and with *C. campestris* attached (labeled with +Cc). (P) Cortex lignin area percentage in different cultivars. Data presented are assessed using multiple comparisons with Dunnett’s test. “*”: p-values < 0.05, “**”: p-values < 0.01. (Q) Cell death area percentage in different cultivars. (P-Q) H1706-Cc, n = 20; H1706+Cc, n = 38; H9775-Cc, n = 19; H9775+Cc, n = 40; H9492-Cc, n = 16; H9492+Cc, n = 38; H9553-Cc, n = 17; H9553+Cc, n = 30. Data were collected at 14 DPA. The data points labeled with grey color indicate the sample that we show in the section picture H-I.

To identify the basis for resistance, we analyzed *C. campestris* attachments on susceptible and resistant lines using anatomy and cell wall-specific staining with Toluidine Blue O and Phloroglucinol-HCl (Liljegren, 2010). Upon challenging these different cultivars with *C. campestris* strands, lignin accumulation in the stem cortex was observed in the resistant H9492 and H9553, but not in the susceptible H9775 and H1706 (Figure 1D – 1O). The resistance mechanism involved local lignification in the stem cortex, creating a barrier to haustorium penetration, and dodder attachment on the resistant cultivars (Figure 1D – 1O). Little to no lignin accumulates in the cortex of both resistant and susceptible cultivars without *Cuscuta* attachment (Figure 1P). In addition, *Cuscuta* attachment sites usually cause some wounding responses and programmed cell death in both resistant and susceptible cultivars (Figure 1Q).

### Identifying the key time point in host-parasite interactions

Changes in the levels of Salicylic Acid and Jasmonic Acid have been reported at 36 to 48 hours after attachment (RUNYON et al., 2010). To capture the earliest responses to dodder parasitism, we performed a time-course RNA-Seq analysis on 0, 1, 2, 3, and 4 DPA (days post attachment) of *C. campestris* on tomatoes H1706 (susceptible). At these stages, the dodder strands were not embedded in the host and could be removed to collect the attached stem area. Maximal transcriptional changes peaked at 4 DPA (Supplemental Figure 1, Supplemental Data Set 1), suggesting that the Differentially Expressed (DE) genes include core genes involved in the early response to *C. campestris* infection. Accordingly, we chose 4 DPA for further gene expression analysis of the resistant and susceptible cultivars.

### Gene expression in the resistant and susceptible host response to *C. campestris*

We challenged the resistant H9492 and H9553, and susceptible H9775 and H1706 with *C. campestris* strands. We collected stem tissues at 4 DPA for RNA-seq and differential gene expression analysis in dodder infested versus uninfested plants. In principal component analysis (PCA) on the transcriptomes of resistant and susceptible cultivars (Supplemental Figure 2), PC1 accounted for 44% of the variation and significantly clustered the data into two separate sets: infested and non-infested samples. However, PCA did not separate different cultivars into distinct genotypic groups. Thus the transcriptional differences in response to *C. campestris* between the resistant and susceptible genotypes likely involve a small number of genes.

Next, we conducted differential gene expression (DGE) analyses by comparing *C. campestris* infested and uninfested host plants using an interaction design model (design model = infested or uninfested condition + genotype + condition: genotype). Based on our communication with the Kraft Heinz Company, both H9492 and H9553 were developed in the same breeding program. However, H9553 is more resistant to *C. campestris* than H9492 based on the relative growth rate results (Figure 1C). Therefore, we suspected that enhanced resistance to *C. campestris* is due to the alterations in key regulatory genes. We selected 113 genes that were differentially expressed (Supplemental Data Set 2) between infested H9775 (susceptible) and infested H9553 (resistant) with adjusted p value cutoff < 0.1 and log2 fold change > 1. Consistent with our observations of lignin accumulation in resistant tomato cultivars upon *C. campestris* infestation (Figure 1), many of these genes are known to be involved in the lignin biosynthetic pathway, including three Laccase genes (LAC4, 5 and 17, Solyc05g052340, Solyc09g010990, Solyc10g085090) and Caffeoyl CoA 3-O-methyltransferase (CCoAOMT, Solyc01g107910) (Figure 2).

**Figure 2.**
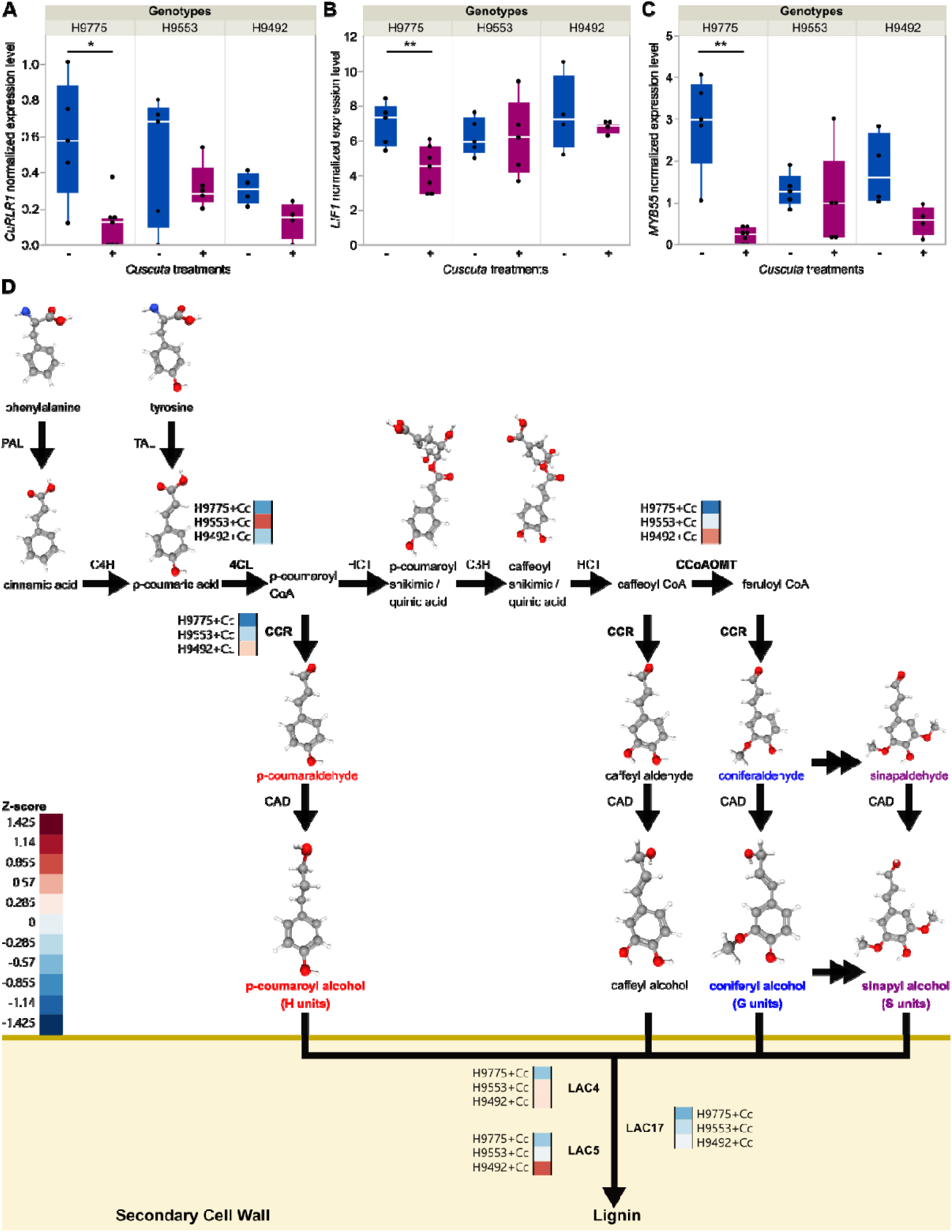
Key candidate genes and lignin biosynthesis genes that display expression changes upon *C. campestris* infestation. (A-C) The normalized expressions levels (CPM, counts per million) of genes in susceptible cultivar H9775 and in resistant hybrid cultivar H9553 and H9492 under *C. campestris* infestation. – and + indicates without or with *C. campestris* infection treatments respectively. Biologically independent replicates: RNA-Seq libraries: H9775-Cc, n = 5; H9775+Cc, n = 7; H9492-Cc, n = 4; H9492+Cc, n = 4; H9553-Cc, n = 5; H9553+Cc, n = 5. Data are assessed using two-tailed t test. “*”: p-values < 0.04, “**”: p-values < 0.01, “***”: p-values < 0.005. (D) The lignin biosynthesis pathway with key enzyme expression levels. Genes that are differentially expressed were selected and the normalized expression values across three cultivars were color coded according to z-score. + Cc indicates with *C. campestris* infection treatments. PAL, phenylalanine ammonia-lyase; C4H, cinnamate 4-hydroxylase; TAL, tyrosine ammonia-lyase; 4CL, 4-coumarate CoA ligase; HCT, hydroxycinnamoyl-CoA shikimate / Quinate hydroxycinnamoyltransferase; C3H, *p*-coumarate 3-hydroxylase; CCoAOMT, caffeoyl-CoA O-methyltransferase; CCR, cinnamoyl-CoA reductase; CAD, cinnamyl alcohol dehydrogenase; LAC, laccase. 3D structure images of phenylalanine, tyrosine, cinnamic acid, p-coumaric acid, p-coumaroyl shikimic acid, caffeoyl shikimic acid, p-coumaraldehyde, p-coumaryl alcohol, caffeyl aldehyde, caffeyl alcohol, coniferaldehyde, coniferyl alcohol, sinapaldehyde, and sinapyl alcohol are from PubChem (National Center for Biotechnology Information, 2021c, b, d, e, f, g, h, i, j, k, l, m, n, a).

To narrow down the potential upstream candidates regulating this lignin-based resistance, we focused on transcription factors (TF - based on gene annotations) as possible key regulators of lignin biosynthesis pathways, and membrane located or cytosolic receptors that may receive signals from *C. campestris*. Using these two criteria, we identified three candidate genes for further study, including a TF related to *AP2*, a *SlMYB55* TF, and a gene encoding an N-terminal coiled-coil nucleotide-binding site leucine-rich repeat protein (*CC-NBS-LRR*) (Figure 2A-C). These three candidate genes share a common expression pattern of significantly reduced expression levels upon *C. campestris* infestation in the susceptible cultivars. However, expression of these three candidate genes remained almost unchanged from uninfested or was only mildly reduced upon *C. campestris* infestation in resistant cultivars. This result suggests that these candidates might play a role in defense against *Cuscuta*, such that when these genes are not repressed during *C. campestris* infestation, the host plants are more resistant to *C. campestris*.

### Functional characterization of candidate genes using virus-induced gene silencing (VIGS) and virus-based gene expression (VGE)

To validate the function of these candidate genes, an ideal method would be generating knockout mutant plants for further study. However, these resistant tomato lines are F1 hybrids in the Heinz background, and the Heinz cultivars are recalcitrant to transformation. Therefore, we used virus-induced gene silencing (VIGS) to knock down our candidate genes in resistant cultivar H9553 to test the functions of our candidate genes. The *C. campestris* plants grown on *AP2*-like, *SlMYB55,* and *CC-NBS-LRR* VIGS knockdown plants have higher survival rates compared with those growing on mock controls (Supplemental Figure 3). Similar phenotypes were also observed in CCoAOMT and LAC gene knockdown plants (Supplemental Figure 3). These results indicate that these candidate genes might play a role in the lignin-based resistance response. Therefore, when these essential genes were knocked down in resistant tomato cultivars, resistant tomato became more susceptible to *C. campestris*, leading to a higher survival rate of the parasite.

To further evaluate if the candidate genes can confer lignification-based resistance in susceptible tomato cultivars, we cloned *GUS, AP2*-like, *SlMYB55*, and *CC-NBS-LRR* genes into Virus-based Gene Expression (VGE) vectors (vector map in Supplemental Figure 4; sequence in Supplemental Data Set 3) for transient overexpression in the susceptible H1706, which has similar expression patterns of these three candidate genes (Supplemental Figure 5A-C). We saw significant GUS expression in the stem around the injection site (Supplemental Figure 5D-F), and lack of lignification due to the process of injection itself (Figure 3A and D). Hence, we used *GUS*-injected plants as our mock controls for VGE experiments. We sectioned and stained injected stems with lignin-specific Phloroglucinol-HCl for lignin detection. VGE with *SlMYB55* and *AP2*-like successfully overexpressed these genes in the first internode near the injection site and induced stem lignification in the susceptible H1706 without *C. campestris* infestation (Figure 3B–C and G). Therefore, we named this *AP2*-like protein *LIF1* (*Lignin Induction Factor 1*) based on its ability to induce lignin biosynthesis in the cortex. These results indicate that *SlMYB55* and *LIF1* might play a role in regulating some of the critical enzymes in the lignin biosynthesis pathway.

**Figure 3.**
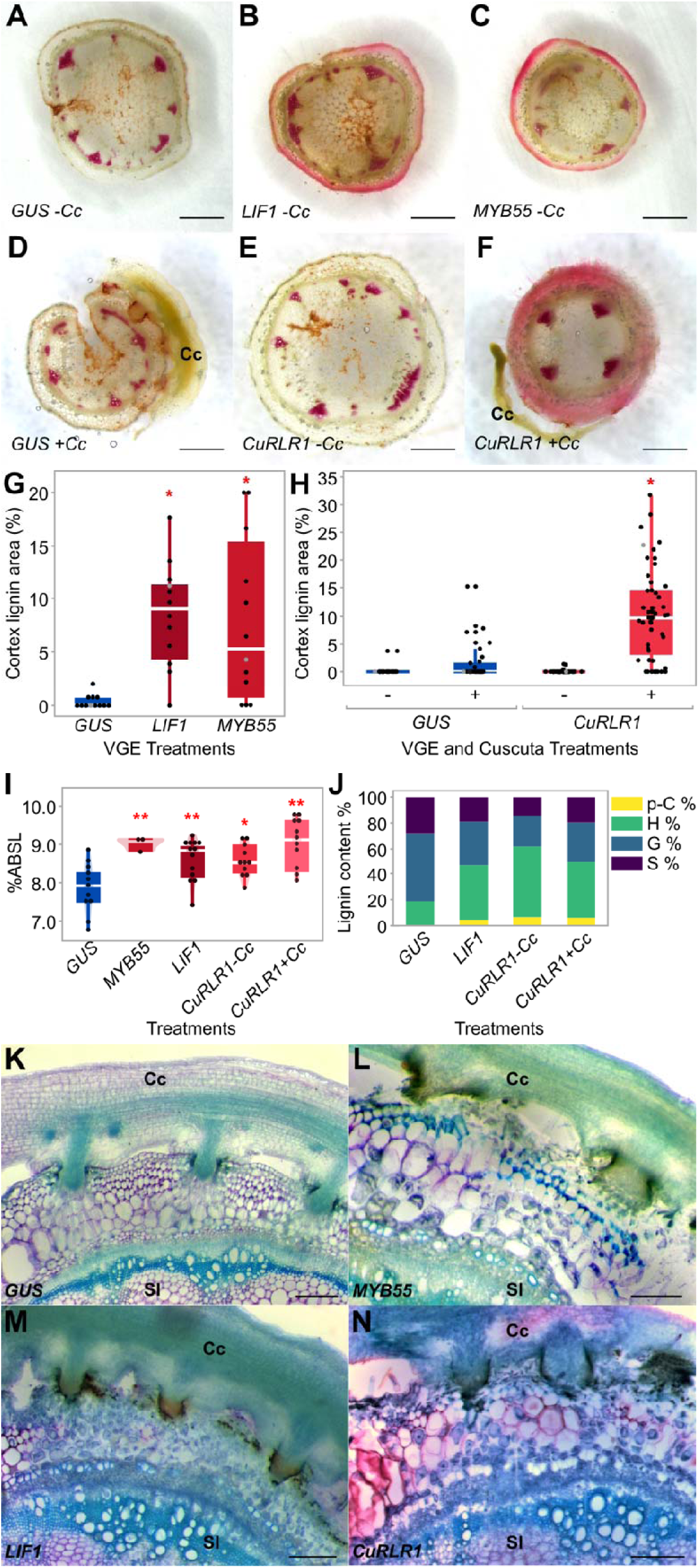
Virus-based Gene Expression (VGE) in tomato H1706. (A-F) ∼300 μm stem sections near injection sites, Phloroglucinol-HCl stains lignin red. VGE of *GUS* (A), *LIF1* (B), *SlMYB55* (C), and *CuRLR1* (E) in stem without *C. campestris*. VGE of *GUS* (D) and *CuRLR1* (F) with *C. campestris*. (G) Cortex lignin area percentage in VGE of *LIF1* and *SlMYB55* (n = 12 each) and (H) *CuRLR1* with and without *C. campestris* (*GUS*-Cc, n = 18; *GUS*+Cc, n =28; *CuRLR1*-Cc, n = 47; *CuRLR1*+Cc, n = 53). (G-H) Data were collected at 7 days post injection (DPI) and 14 days post attachment (DPA). Data are assessed using Dunnett’s test with *GUS*-Cc as negative control. “*”: p-values < 0.01. The data points labeled with grey color indicate the sample that we show in the section picture A-F. (I) Acetyl bromide assay for lignin in VGE stems. Acetyl Bromide Soluble Lignin (ABSL) indicates percent absorbance of soluble lignin. Samples were collected at 7 DPI and 6 DPA. Data are assessed using Dunnett’s test. “*”: p-values < 0.05, “**”: p-values < 0.01. Biological replicates for *GUS*, n = 18; *SlMYB55*, n = 10; *LIF1*, n = 18, *CuRLR1*-Cc, n = 18; *CuRLR1*+Cc, n = 18. Technical replicates for acetyl bromide assay; *GUS*, n = 11; *SlMYB55*, n = 3; *LIF1*, n = 13; *CuRLR1*-Cc, n = 11; *CuRLR1*+Cc, n = 11. (J) PYRO-GC assay for monolignols in *CuRLR1* VGE samples with and without *C. campestris*. Samples were collected at 7 DPI and 6 DPA. Biological replicates collected from first internodes; *GUS*, n = 8; *LIF1*, n = 8, *CuRLR1*-Cc, n = 18; *CuRLR1*+Cc, n = 18. PYRO-GC assay technical replicates; *GUS*, n = 3; *LIF1*, n = 3; *CuRLR1*-Cc, n = 5; *CuRLR1*+Cc, n = 5. (K-N) VGE of *CuRLR1*, *LIF1* and *SlMYB55* induces cortical lignin making H1706 resistant to *C. campestris*. Scale bar, 30 µm. Samples were collected at 7 DPI and 6 DPA. Cc indicates *C. campestris*; Sl indicates *S. lycopersicum*.

In contrast, the H1706 plants with VGE of *CC-NBS-LRR* had no lignin accumulation phenotype and were very similar to those with GUS VGE under no *C. campestris* infestation conditions (Figure 3E). However, previous studies indicated that many genes in the *NBS-LRR* family encode intracellular receptors that detect pathogens and trigger defense signaling (Padmanabhan et al., 2013). Therefore, we suspected that this *CC-NBS-LRR* might play a role in signaling or function as a receptor for signals from *Cuscuta* that are needed to initiate subsequent defense responses. Hence, *C. campestris* infestation treatment might be needed to see the phenotype difference. To validate this hypothesis, we compared the response differences between *Cuscuta* infested and uninfested susceptible H1706 with *CC-NBS-LRR* VGE (Figure 3E – F and H). Intriguingly, our results showed the overexpression of *CC-NBS-LRR* only induced lignification upon *C. campestris* attachment (Figure 3H), and these results suggest that direct or indirect perception of *C. campestris* signals by this *CC-NBS-LRR* leads to lignification-based resistance. Thus, we named this gene *CuRLR1* (*Cuscuta R-gene for Lignin-based Resistance 1*).

On the other hand, we are also aware that lignin is a complex polymer and phloroglucinol-HCl staining is a fast and efficient lignin detection method, but it only detects the cinnamaldehyde end groups of lignin, preferentially staining the G and S-type aldehyde form monolignols (Pomar et al., 2002; Cass et al., 2015). Therefore, we also conducted an acetyl bromide assay to determine total lignin content, including different types of monolignols and lignin precursors. Consistent with the aforementioned anatomical observations, the overexpression of *SlMYB55* and *LIF1* both increased total lignin content compared with GUS mock controls in this assay (Figure 3I). Surprisingly, the overexpression of *CuRLR1* also increased the total lignin content even without *Cuscuta* signals. With *Cuscuta* signals, the total lignin content was much higher in *CuRLR1* overexpressing plants (Figure 3I). This difference indicates that the composition of induced lignin might be different between *CuRLR1* overexpressing plants with and without *Cuscuta* signals.

To further validate this hypothesis, we used high-performance liquid chromatography (HPLC) and pyrolysis gas chromatography/mass spectrometry (Pyrolysis-GC-MS) to analyze the composition of induced lignin. Our HPLC results showed that p-coumarate and trans-ferulate are both increased in *CuRLR1* overexpressed plants, but the samples with *Cuscuta* signals have much higher levels of these two precursors than the samples without *Cuscuta* signals (Supplemental Figure 6). PYRO-GC-MS analysis showed that samples from *CuRLR1* overexpressing plants without *Cuscuta* signals have the larger percentage of H-lignin and the larger concentration of coumarate derivatives compared to VGE of *GUS*, *LIF1*, and *CuRLR1* with *Cuscuta* (Figure 3J). These results show that *CuRLR1* overexpression alone leads to an increase in the upstream steps of the lignin biosynthesis pathway and production of more lignin precursors and H-type monolignols, while adding *Cuscuta* signals may actually up-regulate the final steps in the biosynthesis pathway leading to more G-type and S-type monolignol formation (Figure 3I and J). Since H-lignin and coumarate are not incorporated into lignin as aldehydes, they are not detected by phloroglucinol staining, which explains the difference that we observed between the phloroglucinol staining data and acetyl bromide assay. This phenotype of induced lignin precursors and H-lignin also indicates the *CuRLR1* overexpression alone has turned on the baseline of defense mechanisms. Based on previous studies, H-lignin has been correlated with both stress response as well as defense from pathogen intrusion because this is a form of “defense” lignin that can be generated and deposited more rapidly than G or S lignin (Zhang et al., 2007; Moura et al., 2010; Liu et al., 2018). This baseline of defense mechanisms can then be upgraded upon detecting *Cuscuta* signals, and start accumulating more G-type and S-type monolignols to reinforce a stronger physical boundary.

Eventually, whether or not the overexpression of these candidate genes makes susceptible tomatoes resistant to *C. campestris* is the central question when evaluating potential agricultural applications. Therefore, we transiently overexpressed *SlMYB55*, *LIF1*, and *CuRLR1* first and then attached *C. campestris* strands to test their resistance status. Based on our results, VGE of *SlMYB55*, *LIF1*, and *CuRLR1* with *C. campestris* all induced lignin accumulation in the cortex and blocked haustorium penetration, which made the susceptible tomato cultivar H1706 more resistant to *C. campestris* (Figure 3K – N, Supplemental Figure 7, and Supplemental Data Set 4).

### Regulatory mechanisms and networks leading to resistance responses

Since both H9492 and H9553 hybrid cultivars arose in the same breeding program, enhanced resistance to dodders observed in these two cultivars is likely due to the presence of some unique sequence polymorphisms in these cultivars. Resistance-specific nucleotide polymorphisms (SNPs) could contribute to the regulation or function of our candidate genes, so we specifically identified SNPs that are common in H9553 and H9492 but different from H9775 (Supplemental Data Set 5). Unexpectedly, there were no resistance-specific SNPs in coding regions of our candidate genes except one SNP located in a *LIF1* exon. This resistance-specific SNP changes 251 Lysine (K, in H1706) to 251 Glutamine (Q, in H9553). However, based on our protein domain prediction using InterProScan (Jones et al., 2014; Mitchell et al., 2018) and protein structure analysis using Phyre2 (Kelley et al., 2015), this amino acid replacement is not located in any known protein domains or structures. We also conducted PROVEAN (Protein Variation Effect Analyzer) analysis, and this K251Q variant only has a 0.619 PROVEAN score, indicating that it is a neutral variant. Thus, we conclude that there are no resistance-specific SNP in coding regions of our candidate genes that might contribute to the regulation of resistance.

We, therefore, specifically focused on resistance-specific SNPs in the promoter regions of our candidate genes (Supplemental Data Set 6). Our SNP analysis detected several resistance-specific SNPs in the *LIF1* promoter region, but no resistance-specific SNPs were detected in other candidate gene promoter regions (within 5 kb upstream). One resistance-specific SNP was detected in the *CuRe1* promoter region (outside 5 kb upstream) located at a putative YABBY binding site. However, this SNP is also located 1184bp upstream of *ULP1* (Solyc08g016275) and may regulate expression of this neighboring gene instead of *CuRe1*. Therefore, we focused on the *LIF1* promoter region for further analysis and conducted transcription factor (TF) binding site predictions.

Based on our phylogenetic network analysis (Solís-Lemus et al., 2017) using 500 kb around the *LIF1* resistance-specific SNP enriched region, these SNPs might be introgressed from wild tomato species (likely coming from *S. galapagense* and/or *S. pennellii*, Supplemental Figure 8). One of these resistance-specific SNPs is located right at a WRKY binding W-box cis-element (TTGACY-core motif (Ciolkowski et al., 2008; Chen et al., 2019)) (Supplemental Figure 9 and Supplemental Data Set 7). This SNP is predicted to interrupt WRKY binding, likely leading to *LIF1* expression differences between resistant and susceptible cultivars upon *C. campestris* attachment. Hence, we were also interested in searching for potential *WRKY* TFs in our selected gene lists.

To understand the relationships between the three candidate genes and their targets, identify the potential *WRKY* regulator, and also investigate whether these newly discovered candidate genes connect with previously identified *CuRe1*-mediated resistance responses (Hegenauer et al., 2016), we conducted DGE analysis with ANOVA and selected 10939 differentially expressed genes (DEG) with FDR less than 0.1 (Supplemental Data Set 8). Next, we used Barnes-Hut t-distributed stochastic neighbor embedding (BH-SNE) to generate gene clusters using RSMod (a pipeline developed by us, script included in code availability) (Ranjan et al., 2016). In this analysis 5941 DEGs are clustered into 48 groups based on their gene expression patterns and 4998 DEGs are in the noise group. Among the 48 gene clusters generated (Supplemental Data Set 9), five clusters were selected based on their GO (Gene Ontology) enrichment terms and the candidate genes they included (Figure 4A-B and Supplemental Data Set 9). The GO term of cluster 39 is “DNA binding”, which includes potentially key TFs, like *MYB55*. The GO term of cluster 11 is “lignin biosynthetic and catabolic process”, which encapsulates the observed resistance phenotypes, and includes Caffeoyl CoA 3-O-methyltransferase (*CCOMT*) and three Laccase (*Lac*) genes identified in our model-based approach (Supplemental Data Set 2). Cluster 23 includes a *Cinnamoyl-CoA reductase* gene (Solyc03g097170, *CCR*) and is enriched in the “xyloglucan:xyloglucosyl transferase activity” GO term, which may indicate potential cell wall modifications. “Response to biotic stimulus” is the GO term enriched in cluster 17, which also includes the previously identified Cuscuta receptor, *CuRe1*.

**Figure 4.**
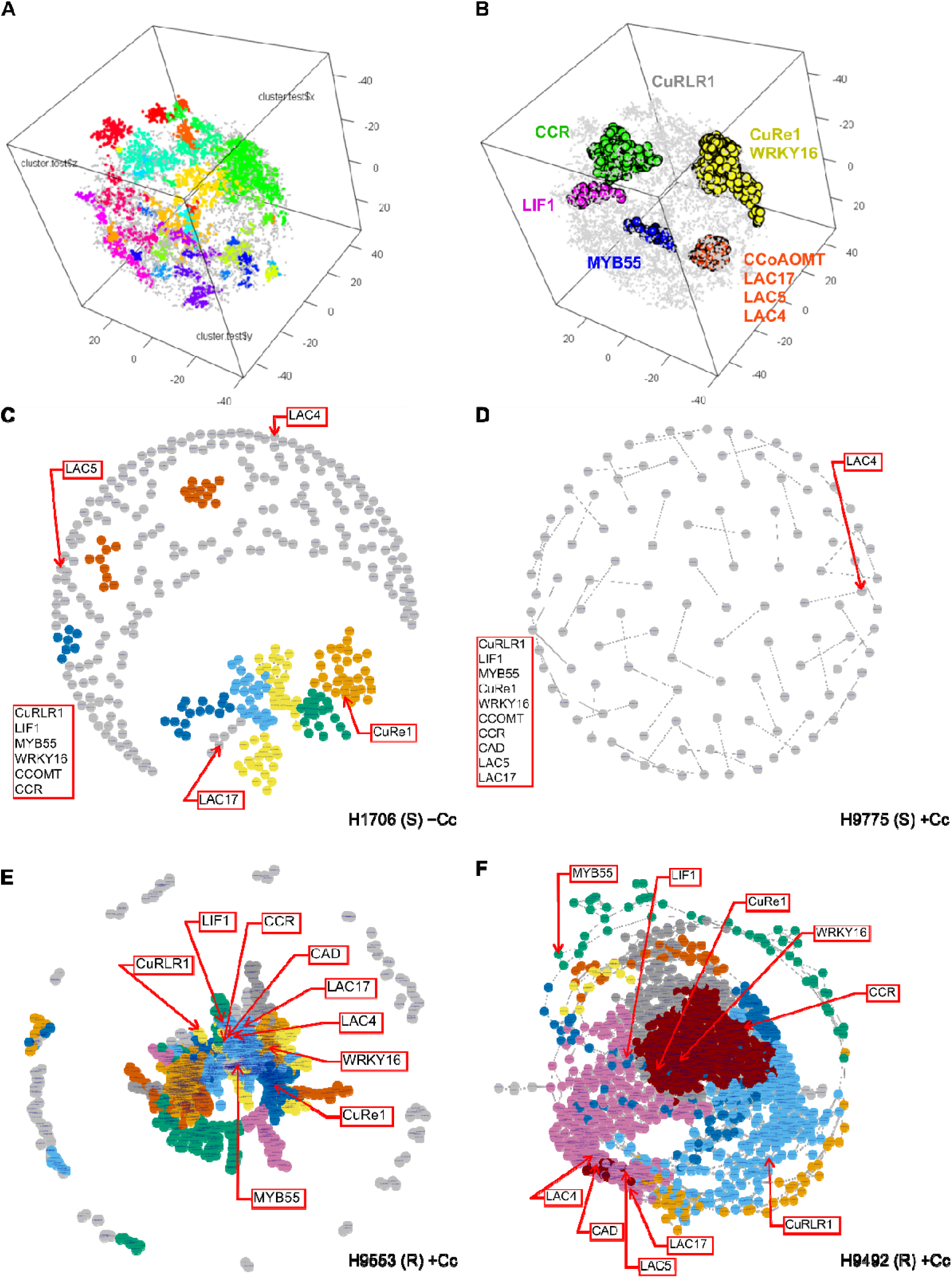
Barnes-Hut t-distributed stochastic neighbor embedding (BH-SNE) generated gene clusters and gene co-expression network (GCN) analysis. (A) BH-SNE generated gene clusters based on their gene expression patterns. Total cluster (module) number was 48. (B) The candidate genes that are included in the clusters are labeled with their corresponding colors. The selected gene clusters for GCN are labeled in yellow (CuRe1 and WRKY16 cluster, cluster 17), red (CCOMT and LAC cluster, cluster 11), blue (MYB55 cluster, cluster 39), pink (LIF1 cluster, cluster 46), green (CCR cluster, cluster 23) colors. *CuRLR1* is in the noise cluster, and is labeled in grey color. Parameters used in this analysis: perplexity (perp) = 20, lying = 250, cutoff = 20, seed = 2. (C-F) Gene co-expression networks (GCNs) of four different Heinz susceptible and resistant cultivars upon *C. campestris* treatments. Based on BH-SNE analysis, 1676 genes in cluster 11, 17, 23, 39, 46 and CuRLR1 were selected for building GCNs. +Cc indicates with *C. campestris* infection treatments. The genes that are listed at the left of the GCN and not labeled in the network are the genes that have no coexpression connections with all the other genes in the list.

Additionally, with comprehensive RNA-Seq clustering and gene-coexpression analysis results, we also noticed *SlWRKY16* (Solyc07g056280) is always clustered with *CuRe1*. *SlWRKY16* was highly upregulated at 4 DPA in all four Heinz cultivars, an expression pattern similar to that for *CuRe1* (Figure 5A-B). Host tissues surrounding haustoria from the tomato M82 cultivar also show upregulated expression of *SlWRKY16* at 4 DPA in our time-course data with FDR < 0.1 and real-time qPCR data (Supplemental Figure 10). Thus, *SlWRKY16* is a commonly upregulated host response gene across different cultivars and may play an important role in the transduction of *C. campestris* signals upon host attachment. Furthermore, one of the resistance-specific SNPs in the *LIF1* promoter region mentioned above, is located at a WRKY transcription factor W-box (TTGACY-core motif) binding site, which is also the predicted *SlWRKY16* binding site based on homologous genes in the phylogenetic tree of the WRKY domain at the Plant Transcription Factor Database (Jin et al., 2016). Taking all these criteria together, we included *SlWRKY16* in our candidate genes for further analysis.

**Figure 5.**
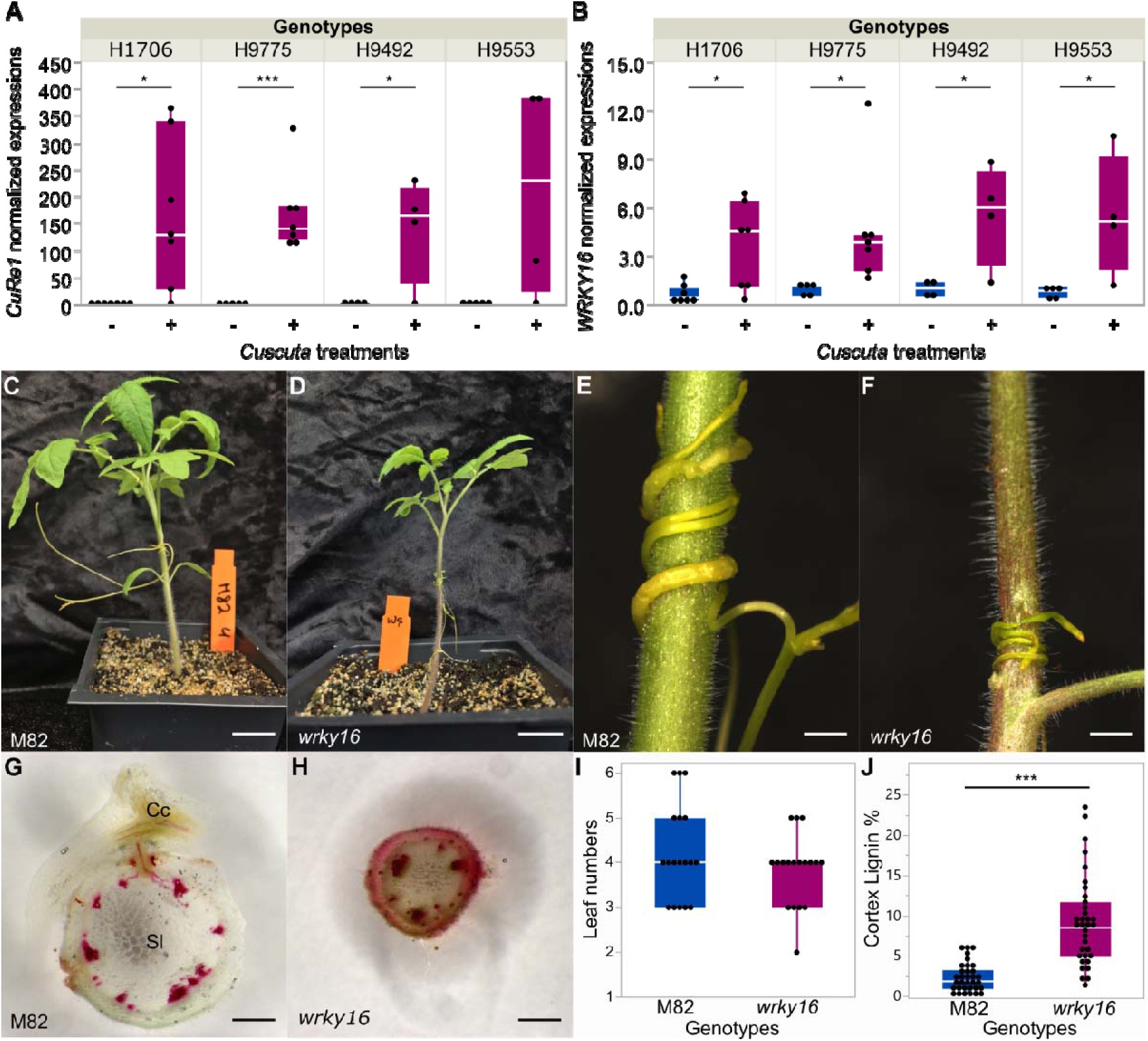
The role of *SlWRKY16* in *Cuscuta* resistance using CRISPR/Cas9 gene knockouts. (A-B) Normalized *CuRel* and *SlWRKY16* expression level from RNA-Seq data (CPM, counts per million) in different Heinz cultivars with/without *Cuscuta* treatments. Biologically independent replicates: RNA-Seq libraries, H1706-Cc, n = 7; H1706+Cc, n = 7; H9775-Cc, n = 5; H9775+Cc, n = 7; H9492-Cc, n = 5; H9492+Cc, n = 4; H9553-Cc, n = 5; H9553+Cc, n = 5. Data are assessed using one-tailed t test. “*”: p-values < 0.05, “**”: p-values < 0.01, “***”: p-values < 0.005. (C-J) Samples and data were collected at 7 DPA. (C-D) overall phenotype comparison between M82 and homozygous *SlWRKY16* CRISPR lines (*wrky16*). Scale bar, 2 cm. (E-F) *C. campestris* growing on M82 and *wrky16*. (G-H) ∼300 μm hand sections of M82 and *wrky16* stems near *Cuscuta* attachment site stained with Phloroglucinol-HCl. Lignin is stained red. Cc indicates *C. campestris.* (I) Leaf number of *wrky16* and M82. Biological replicates, n = 18 for each. (J) Cortex lignin area percentage in M82 and *wrky16* stems. Data presented are assessed using student’s t-test. “***” indicates p-value is less than 0.001. Replicates: M82, n = 33; *wrky16*, n = 34.

We focused on these 1676 genes in clusters 11, 17, 23, 39, 46 and included *CuRLR1* (Figure 4A-B and Supplemental Data Set 9) to construct gene co-expression networks (GCNs) for different treatments and cultivars to identify central hub genes (Figure 4, Supplemental Figure 11) (script included in code availability). Interestingly, *CuRLR1*, *SlWRKY16* and *CuRe1* had few connections or almost no connection with other genes in the GCN in susceptible cultivars with *Cuscuta* attachments (Figure 4C-D). On the other hand, *CuRe1* and *SlWRKY16* became central hub genes in resistant cultivars only upon *C. campestris* attachments and connected with *CuRLR1* (Figure 4E-F). However, based on our DNA-Seq analysis, we cannot detect any resistance-specific SNPs in the promoter regions or coding regions of *CuRe1* and *CuRLR1*, and *SlWRKY16* (Supplemental Data Set 5). This result indicates that the differential expression of *CuRe1* and *CuRLR*1, and *SlWRKY16* may be controlled by trans-regulatory factors or protein interactions. Based on our GCN analysis and DNA-Seq analysis results, we propose that all four Heinz tomato cultivars have *CuRe1*, *CuRLR1*, and *SlWRKY16*. Among them, *SlWRKY16* is a key factor in the transduction of *C. campestris* signals upon attachment of the parasite to the host. However, the differential expression patterns upon *C. campestris* attack and the diverse regulatory connections of these three genes determine whether resistance responses are triggered in these Heinz cultivars or not.

### Functional characterization of *SlWRKY16* by CRISPR/Cas9 knockouts and VGE

Since *SlWRKY16* exists in all Heinz resistant and susceptible tomatoes and in the M82 tomato cultivar, we bypassed the transformation limitation in Heinz tomatoes and generated mutant tomato plants in the M82 background for further analysis. To validate the function of *SlWRKY16* and its role in lignification-based resistance, we produced stable *SlWRKY16* edited M82 lines using the CRISPR/Cas9 targeted gene knockout system (Pan et al., 2016). Our homozygous null mutants were generally smaller than M82 wild type (Figure 5C and D) even though both *wrky16* and M82 wild type show the same developmental progression (Figure 5I). Intriguingly, *wrky16* plants are more resistant to *C. campestris* than M82 wild type (Figure 5E – H). Using Phloroglucinol staining, we noticed that homozygous *wrky16* lines continuously produce cortical lignin, which forms a physical boundary and provides a strong resistance to *C. campestris* attachment compared to M82 wild type (Figure 5E – H and J). However, the phenotype of continuously accumulating cortical lignin likely also limits cell growth and leads to the stunted growth phenotype in *wrky16* plants. These results indicate that *SlWRKY16* may function as a negative regulator of the lignin-based resistance response.

The hypothesis that *SlWRKY16* may play a role in the lignin-based resistance response also incorporates our previous SNP analysis and transcription factor binding site prediction results in the *LIF1* promoter region (Supplemental Figure 9). We proposed that the resistance-specific SNP located at a WRKY binding site in the *LIF1* promoter region could interrupt SlWRKY16 protein binding, leading to *LIF1* expression differences between resistant and susceptible cultivars upon *C. campestris* attachment. Therefore, we conducted real-time qPCR to determine the expression levels of *LIF1* in both susceptible M82 wild type and resistant *wrky16* tomatoes (M82 background). We observed a mild increase in *LIF1* expression in *wrky16* tomatoes compared to M82 tomatoes (Supplemental Figure 12A). Considering LIF1 is an AP2/B3-like transcription factor, any elevation in *LIF1* expression could potentially lead to large differences in the downstream gene expression pathways.

To evaluate the interaction between *SlWRKY16* and the other three candidate genes, we transiently overexpressed *LIF1*, *SlMYB55*, *CuRLR1,* and *GUS* controls in the susceptible H1706, M82 wild type, and resistant *wrky16* tomatoes. In the *GUS* transient overexpression control group, we observed that *wrky16* plants accumulate much more lignin than H1706 and M82 wild type as expected (Supplemental Figure 12B). Overexpression of *LIF1* induced more lignification in H1706, M82, and *wrky16* plants. This result shows additive effects of loss of *SlWRKY16* function and overexpression of *LIF1* in lignification responses (Supplemental Figure 12B), suggesting that *SlWRKY16* may not only regulate *LIF1* expression at the transcriptional level, but also may regulate LIF1 protein function by other mechanisms. Also, overexpression of *MYB55* induced more lignification in H1706 and *wrky16* plants (M82 background) but not in M82 (Supplemental Figure 12B), indicating that the loss of *SlWRKY16* function in M82 allows more lignin accumulation upon *MYB55* overexpression. This result also suggests subtle differences in resistant response between cultivars and that *SlWRKY16* might act upstream of *MYB55*, but more details remain to be elucidated in future research.

On the other hand, overexpression of *CuRLR1* with *C. campestris* infection was able to induce more lignification in M82, but not in *wrky16* tomatoes (Supplemental Figure 13). This epistatic phenotype suggests that either *CuRLR1* and *SlWRKY16* are in the same pathway with WRKY16 downstream of CuRLR1, or that *CuRLR1* and *SlWRKY16* are in two independent pathway that may influence each other. This hypothesis matches with the gene coexpression networks we built, which show that *CuRLR1* and *SlWRKY16* are peripherally positioned in resistant cultivars in the *Cuscuta* treated condition, with multiple layers of genes connecting them. In order to elucidate other layers of regulation between these genes, we conducted protein-protein interaction investigations.

### Subcellular localization and interactions between the candidate proteins

One described mechanism for triggering innate immunity following TMV infection in tobacco involved interaction and subsequent nuclear localization of the SPL6 TF with the TIR-NBS-LRR receptor (Padmanabhan et al., 2013; Padmanabhan and Dinesh-Kumar, 2014). Therefore, we investigated the potential interactions between our candidates and their protein subcellular localization to uncover potential regulatory mechanisms. Based on our results using translational GFP fusions, LIF1 and SlWRKY16 are located mainly in the nucleus (Figure 6A), while *CuRLR1* is located in both the nucleus and the cytosol. Bimolecular fluorescence complementation (BiFC) experiments with split YFP using transient infiltration in *N. benthamiana* leaves show that the LIF1 and SlWRKY16 proteins interact and get localized to the cytoplasm (Figure 6B). Interactions between other combinations, CuRLR1-LIF1, CuRLR1-SlWRKY16, or CuRLR1-CuRe1, were not detected in our experiments (Figure 6B).

**Figure 6.**
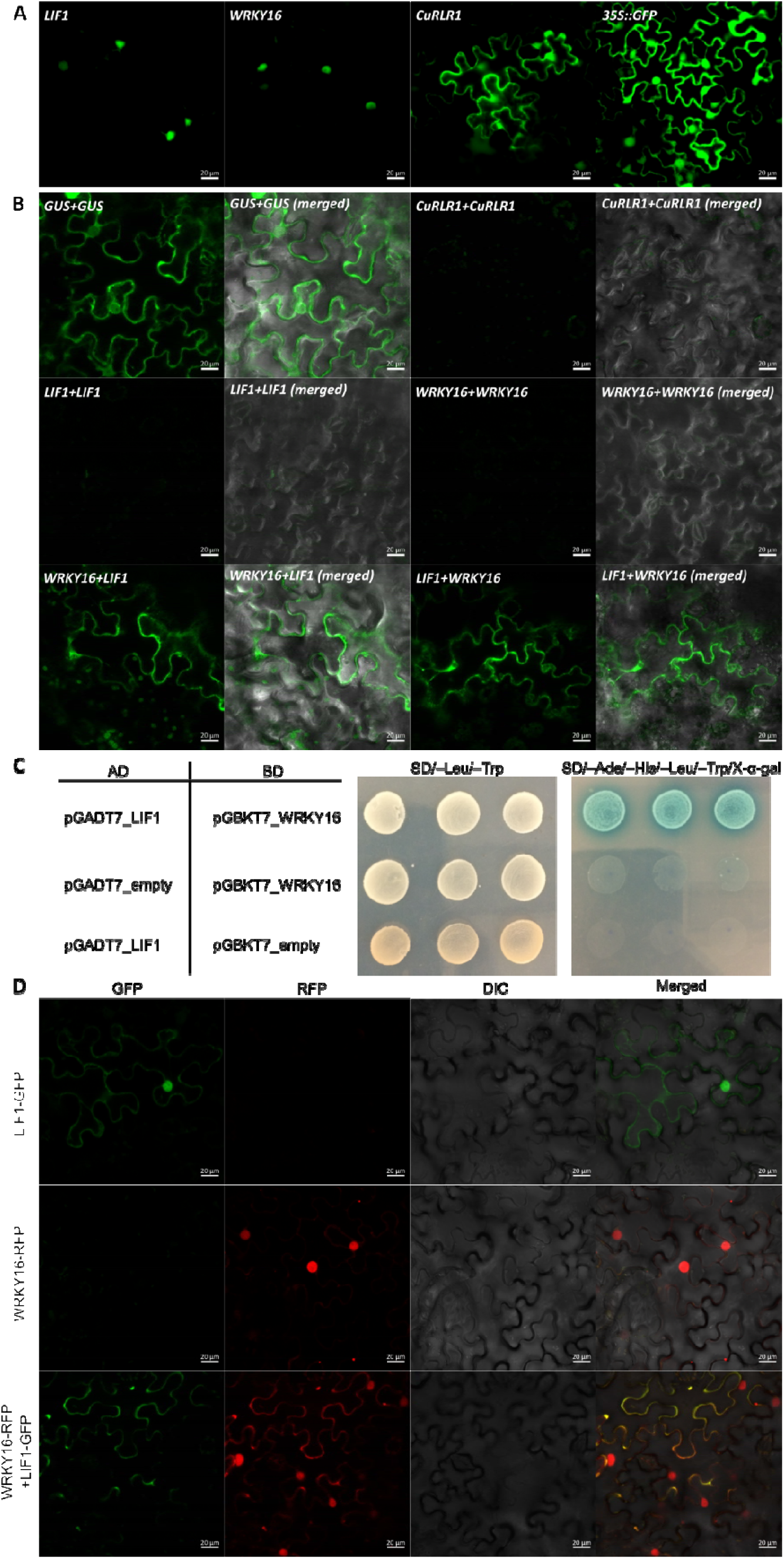
Subcellular localization of candidate genes and protein-protein interactions. (A) Subcellular localizations of LIF1, SlWRKY16, and CuRLR1 proteins. (B) Verification of protein-protein interactions and locations of SlWRKY16 and LIF1 by Bimolecular Fluorescence Complementation (BiFC). The gene with *cCitrine* fusion is listed before the “+” sign and the gene with *nCitrine* fusion is listed after the “+” sign. (C) Yeast two-hybrid (Y2H) results for interaction between LIF1 and SlWRKY16. The plasmids with GAL4 activation domain (AD) and GAL4 DNA binding domain (BD) were co-transformed to yeast AH109 competent cells. Transformed yeast cells were screened on SD/-Leu/-Trp medium plates to select successful co-transformants and then assayed by culturing on high-stringency SD/–Ade/–His/–Leu/–Trp medium plates with 40 μg/ml X-α-Gal. The positive protein-protein interactions between LIF1 and SlWRKY16 are indicated by growth on SD/–Ade/–His/–Leu/–Trp/X-α-Gal medium plates and blue colony color. (D) Co-expression of fusion protein LIF1-GFP and SlWRKY16-RFP to observe subcellular localizations. Yellow color in the merged panel indicates that GFP and RFP signals are overlapped.

To further validate the interaction between LIF1 and SlWRKY16 proteins, we used the GAL4 yeast two-hybrid (Y2H) assays with the yeast (*Saccharomyces cerevisiae*) strain AH109 for examination. Growth on SD/–Ade/–His/–Leu/–Trp/X-α-Gal medium plates and blue colony color confirmed that LIF1 indeed interacted with SlWRKY16 (Figure 6C). To verify the interaction between LIF1 and SlWRKY16 proteins and their subcellular localizations when they interact with each other, we also co-expressed the fusion proteins LIF1-GFP and SlWRKY16-RFP. We found that GFP and RFP signals are located mainly in the nucleus when we only overexpress LIF1-GFP or SlWRKY16-RFP in separate *N. benthamiana* leaves (Figure 6D). However, when we co-express LIF1-GFP and SlWRKY16-RFP in the same leaves, GFP and RFP signals mostly overlap in the cytoplasm (Figure 6D). These results not only further confirm that LIF1 and SlWRKY16 proteins may interact with each other and become cytosol localized, but also validate our hypothesis that SlWRKY16 can regulate LIF1 expression at both the transcriptional and protein interaction levels.

### Analysis of the *Cuscuta* signal using *Cuscuta* extract injections

To further discern the nature of the major signals that trigger lignification-based resistance, we injected the first internode of the resistant H9553 with *Cuscuta* extracts subjected to different treatments (Supplemental Figure 14). Untreated or filtered *Cuscuta* extract injections induced the accumulation of lignin in the cortex region (Supplemental Figure 14B-C). On the other hand, alteration of *Cuscuta* extract pH from 5.8 to 9 abolished lignin accumulation (Supplemental Figure 14D-E), suggesting either instability or sequestration of the *Cuscuta* signaling molecules in alkaline conditions. In addition, heat-treated extract and proteases-treated extract could not trigger the lignification response (Supplemental Figure 14F-J). Moreover, *Cuscuta* extract injections also induced lignin accumulation in H1706 with VGE overexpressing *CuRLR1*, but not in H1706 with GUS VGE (Supplemental Figure 15). This result indicates that *CuRLR1* may be able to either sense some unknown factors in *Cuscuta* extract or some part of the response to these factors and lead to lignin-based resistant responses. Furthermore, filtration of extracts through devices with different molecular weight cutoffs indicates that fractions smaller than 30kD cannot trigger strong lignification response (Supplemental Figure 16). Thus, the active *Cuscuta* signal for induction of lignin-based resistance is larger than 30kD but smaller than 100kD, and distinct from the previously identified *Cuscuta* signal that binds *CuRe1* (Hegenauer et al., 2016; Hegenauer et al., 2020).

## Discussion

*Cuscuta* spp. cause massive losses in infested tomato fields in the United States, so understanding the resistance mechanism of these specific Heinz tomatoes will provide the potential of developing crop protection systems. Notably, previous studies indicate that different *Cuscuta* species can have diverse host-parasite interactions with the same host species (Ranjan et al., 2014; Kaiser et al., 2015; Hegenauer et al., 2016). For example, although cultivated tomatoes (*S. lycopersicum*) are generally resistant to *Cuscuta reflexa* (Sahm et al., 1995; Hegenauer et al., 2016), most domesticated tomato cultivars are susceptible to *C. campestris*. Therefore, using the Heinz tomato cultivars that have been bred for resistance to dodders helped us understand the multilayered resistance mechanisms to *Cuscuta* spp. and how this might aid in developing parasitic plant-resistant crops. This study reveals the underlying resistance mechanism is a lignin-based resistance response in these Heinz resistant tomato cultivars.

Lignin is a complex phenolic polymer, which is generated from three major monolignols, paracoumaryl alcohol, coniferyl alcohol, and sinapyl alcohol, using covalent crosslinks formed via free radical polymerization (Ferrer et al., 2008). Accumulation of lignin in plant stems or roots has been shown to reinforce plant resistance to invading herbivores, parasites and pathogens (Reimers and Leach, 1991; Gayoso et al., 2010; Taheri and Tarighi, 2012; War et al., 2012; Dhakshinamoorthy et al., 2014; Kumari et al., 2016; Zhang et al., 2019). Lignification at the host-parasite interface in roots has been reported in plants that are resistant to root parasitic plants (Goldwasser et al., 1999; PÉRez-De-Luque et al., 2005; CAMERON et al., 2006; Lozano-Baena et al., 2007). However, for stem parasitic plants, most research has focused on hypersensitive response or necrosis as the major mechanisms for host plant defense (LANE et al., 1993; Hegenauer et al., 2016; Su et al., 2020). One previous report of incompatible reactions between tomato plants and *Cuscuta reflexa* characterized by a visible brownish plaque at infection sites, suggested this might be due to suberized or lignified cell walls (Sahm et al., 1995). Here, we first identified a strong lignin-based resistance response toward *C. campestris* attack in these specific Heinz tomato cultivars, adding another layer on the previous reported hypersensitive-type response mechanism.

This lignin-based resistance response is regulated by three key genes, *LIF1*, *SlMYB55*, and *CuRLR1*. Of these, *CuRLR1* responded to unknown *Cuscuta* signals and further reinforced lignin deposition in the resistant cultivars. The *Cuscuta* signals that trigger the lignin-based defense responses appear to be large heat-sensitive proteins (30 kDa - 100 kDa, Supplemental Figure 16), and distinct from the previously identified small *Cuscuta* signal 11 kDa glycine-rich protein (GRP) or its minimal peptide epitope Crip21 (Hegenauer et al., 2020) that is recognized by *CuRe1* (Hegenauer et al., 2016). It would be of interest to investigate interactions between these potential *Cuscuta* signals or effectors that interact with the two different *Cuscuta* receptors.

In conclusion, we propose a new multilayered model for *Cuscuta* resistance response in tomato (Figure 7). *CuRLR1* is a cytosolic factor, which either receives large signaling molecules from *C. campestris* as a receptor, or may be a novel factor which plays a role in downstream signal transduction upon sensing *Cuscuta* signals, and triggers a lignin-based resistance response (Figure 7, red-labeled arrow). Based on previous studies, NBS-LRRs are usually located in the cytoplasm and nucleus and likely to recognize pathogen effectors to induce effector-triggered immunity (ETI) (Dodds and Rathjen, 2010). This matches where we observed CuRLR1 subcellular localization (Figure 6) and might also explain why the *Cuscuta* signals that trigger the lignin-based defense responses are in a different size range compared with the previously identified *Cuscuta* signal. Our research results shed light on a potential ETI pathway in parasitic plant resistance and provides the foundation for future studies into how these various layers of resistance connect.

**Figure 7.**
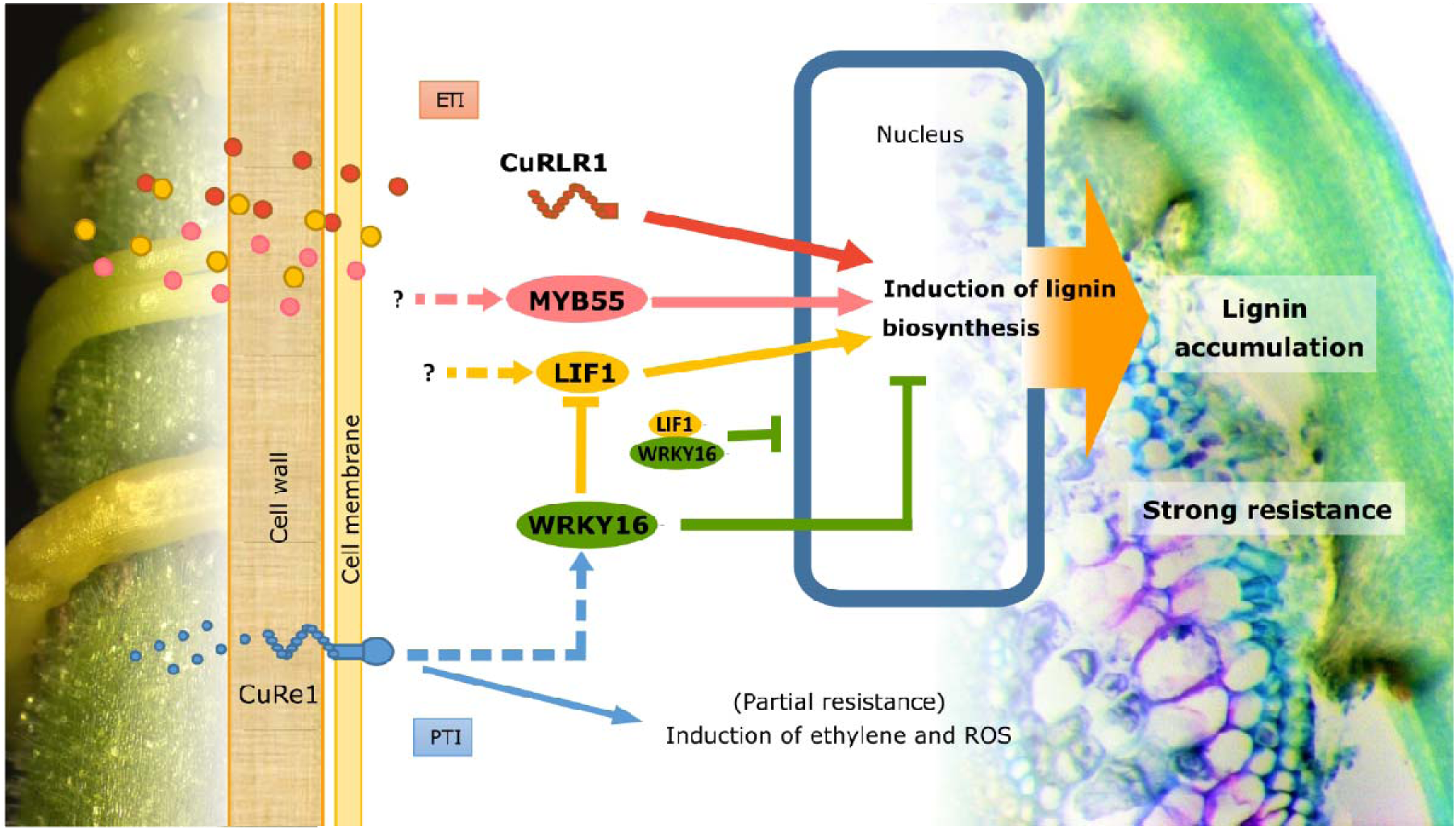
Model of *C. campestris* resistance response in tomato cultivars. Red-labeled pathway: newly identified cytosolic *CuRLR1*, which may receive large signaling molecules from *C. campestris* or play a role in signal transduction upon *Cuscuta* perception. This triggers downstream signal transduction and induces a lignin-based resistance response. This resistant response may be an effector-triggered immunity (ETI). Pink and yellow-labeled pathway: *SlMYB55* and *LIF1* function as positive regulators in the lignin biosynthesis pathway. Yellow and green-labeled pathway: *SlWRKY16* and *LIF1* mediated lignin-based resistant responses and with a potential connection to *CuRe1*. Blue-labeled pathway: previously identified *CuRe1* mediated PAMP/MAMP-triggered immunity (PTI/MTI) pathway.

In our model, *SlMYB55* and *LIF1* were placed as positive regulators in the lignin biosynthesis pathway (Figure 7, pink and yellow-labeled arrows) because transient overexpression of *SlMYB55* and *LIF1* induced lignin accumulation in the cortex (Figure 3). Other yet undiscovered *Cuscuta* receptors or factors may induce *SlMYB55* and *LIF1* expression upon *Cuscuta* attachment. On the other hand, *wrky16* plants showed lignin accumulation and stronger resistance to *Cuscuta*, suggesting that *SlWRKY16* is a negative regulator of this lignin-based resistance pathway (Figure 7, green-labeled arrow). Based on our DNA-Seq, BiFC, and subcellular localization data (Figure 6, Supplemental Figure 9), we propose that *SlWRKY16* regulates the function of *LIF1* by a combination of inhibition of *LIF1* transcription and physical capture of LIF1 proteins to block their entry into the nucleus (Figure 7, yellow and green-labeled arrows). *CuRe1* is reported to mediate PAMP/MAMP-triggered immunity (PTI/MTI) (Hegenauer et al., 2016) (Figure 7, blue-labeled arrow). GCN analysis indicates a coexpression connection between *CuRe1* and *SlWRKY16* (Figure 4A-D). *CuRe1* and *SlWRKY16* both became central hub genes in resistant cultivars upon *Cuscuta* attachments (Figure 4D), suggesting the hypothesis that *SlWRKY16* may act downstream of *CuRe1* (Figure 7, blue-labeled arrow). Thus, we envision crosstalk between different resistance pathways that may be triggered together to enhance host defense responses.

We conclude from our work that the resistance in these specific Heinz tomato cultivars relies on a lignin-based response. The systemic investigation of this resistance response in tomato plants toward the stem parasitic plant *C. campestris* provides potential implications for enhancing crop resistance to parasitic plants. Interestingly, none of the early-step lignin biosynthetic genes, like PAL, C4H, 4CL, were in the model-based differentially expressed gene list. Changing the early steps in lignin biosynthetic genes can also change the phenylpropanoid pathway for the biosynthesis of anthocyanins. This further confirms that lignin biosynthesis is specifically triggered in the Heinz resistant cultivars. Notably, overexpression of the CuRLR1 protein induced upregulation of lignin precursors, but extensive lignin accumulation was only triggered by *Cuscuta* extracts. Introducing CuRLR1 protein could provide resistance to *C. campestris* without triggering the crop to continuously spend a lot of resources producing a large amount of cortical lignin with associated stunted growth. The identification of *CuRLR1* might provide a path forward to introduce resistance into other important crops that are also attacked by *C. campestris*. In summary, *CuRLR1*, *SlWRKY16*, *LIF1*, and *SlMYB55* regulate a lignin-based response in the tomato stem cortex, which prevents *C. campestris* strands from parasitizing these resistant Heinz cultivars.

## Materials and Methods

### Plant materials used in the study

We obtained four different cultivars from Dr. Rich Ozminkowski at HeinzSeed, including the Heinz hybrid cultivars 9492 and 9553 (H9492 and H9553), and the related susceptible Heinz hybrid cultivar 9775 (H9775) and the sequenced susceptible Heinz cultivar 1706 (H1706). Our *Cuscuta* was originally collected from tomato field in California, and we obtained seeds from W. Thomas Lanini. This *Cuscuta* was previously identified as *Cuscuta pentagona* (Yaakov et al., 2001), which is a closely related species to *Cuscuta campestris* (Costea et al., 2015b). To clear up the confusion, we extract DNA to verify species by molecular phylogenetics. Based on phylogenetic analysis of plastid trnL-F, rbcL sequences, and nrITS, nrLSU sequences (Stefanović et al., 2007; García et al., 2014; Costea et al., 2015b), we confirmed that our experimental species is *Cuscuta campestris* (Supplemental Figure 17-21). According to our results, our *Cuscuta campestris* isolate is most similar to *Cuscuta campestris* 201 voucher Rose 46281 WTU from USA, CA (Supplemental Figure 21) that is published by Costea *et al*. in 2015 (Costea et al., 2015a).

### Histology and Cell Wall-Specific Staining

For preparing the sections at the *C. campestris* attachment area and Agroinjection sites on tomato stems, we hand-sectioned plants at 200 to 500 μm thickness using razor blades and kept these sections in 4°C water before staining. For preparing the sections of haustoria attached to host, we fixed samples in 7% Plant Tissue Culture Agar and used Lancer Vibratome Series 1000 to prepare 100 μm sections and kept these sections in 4°C water.

For Phloroglucinol-HCl Staining, we followed the published protocols (Liljegren, 2010; Pradhan Mitra and Loqué, 2014) with some modifications. To prevent plasmolysis during staining, we added an ethanol dehydration process before staining, which is described as follows: we removed the water and then immersed sections in cold 30% ethanol and then 60% ethanol for 5 minutes each. We prepared phloroglucinol-HCl stain (Ph-HCl) or Wiesner stain by preparing a 2:1 mixture of 100% EtOH and concentrated HCl and dissolving powdered phloroglucinol into this solution at a final concentration of 3% w/v. After removing the 60% ethanol, we added phloroglucinol-HCl solution dropwise to the Petri dishes, and let the sections sit in the stain for 5 minutes. The lignified areas of the sections stain bright red within 30 seconds of immersion in the stain. After removing the phloroglucinol-HCl and adding 60% ethanol back, we imaged the sections in the petri dish on a white background using a Zeiss SteREO Discovery, V12 microscope and Nikon Eclipse E600 microscope.

For Toluidine Blue O Staining, we used a published protocol with some modifications (O’Brien et al., 1964). We immersed the sections in the stain for 30 seconds, and then washed with water three times for 30 seconds each. After removing the agar from around the sections using forceps, we mounted the sections with water on a slide and imaged using a Nikon Eclipse E600 microscope.

### Image analysis of stem and haustorium sections

To quantify the lignin content of each section, we analyzed images using the image processing software ImageJ (Schneider et al., 2012). We added a Gaussian blur with a sigma radius of 2.00 to reduce image noise. We set the color space of the image to L*a*b* to generate histograms that measure lightness, green-red contrast, and blue-yellow contrast of the image. We adjusted the lightness filter to allow histogram coordinates ranging from zero to the peak of the image histogram, and the green-red filter to allow from the histogram peak to 255, and the blue-yellow filter to allow all histogram coordinates. These coordinates filter for red areas on the image, corresponding to lignified areas in the stem sections. We measured the total area of lignification, then selected areas corresponding to the lignified xylem of the stem and measured this area. We subtracted the xylem area from the total lignin area to calculate the cortex lignin area.

### DNA-seq library construction for resistant and susceptible Heinz cultivars

DNA was extracted from the leaves of three weeks old seedlings using GeneJET Plant Genomic DNA Purification Mini Kit (Thermo Scientific, Waltham, MA, USA) DNA-Seq libraries were prepared using an in-house protocol modified from Breath Adapter Directional sequencing (BrAD-seq) (Townsley et al., 2015). First, 5 µg of genomic DNA was fragmented using a Covaris E220 (Covaris, Inc. Woburn, MA, USA) with the following settings: Peak Incident Power (W) 140; Duty Factor 10%; Cycles per Burst 200 and Treatment Time (s) 90 to obtain an average fragment size of 400 base pairs. Next, the fragmented DNA was end-repaired and A-tailed in a single reaction using DNA End Repair Mix and Taq DNA polymerase (New England Biolabs). Y-type adapters were ligated and an enrichment PCR was performed with as in BrAD-seq (Townsley et al., 2015) using 7 cycles. Individual libraries were quantified by PCR and pooled to equaled amounts. After a final library cleanup with AMPure beads (Beckman Coulter, Brea, CA, USA), DNA-seq libraries were sequenced at the California Institute for Quantitative Biosciences (QB3) at the University of California, Berkeley using the HiSeq 4000 platform at 150 Paired Read (PR). (Illumina Inc. San Diego, CA, USA).

### Resistant and susceptible Heinz cultivar DNA-seq SNP analysis, promoter binding site analysis, and protein domain and structure prediction

For single nucleotide polymorphism (SNP) analysis, we mapped DNA-seq read data to sequenced H1706 tomato genome itag 3.0 to identify SNPs using CLC Genomics Workbench 11 (QIAGEN, https://www.qiagenbioinformatics.com/). Next, we compared the SNPs across the resistant and susceptible tomato cultivars and focus on finding the SNPs that are common in H9553 and H9492 but different from H9775. In other words, we focus on the SNPs that exist in resistant cultivars and named these SNPs as “resistant specific SNPs” (Supplemental Data Set 5). Among our 4 candidate genes, *LIF1* was the only gene that has resistant specific SNPs in the promoter region (Supplemental Data Set 6). In order to identify potential introgression regions, we conducted phylogenetic network analysis using the PhyloNetworks package (Solís-Lemus et al., 2017) in the Julia environment on XSEDE (Dahan et al., 2014) with 500 kb of sequence around the LIF1 resistance-specific SNP enriched region (Supplemental Figure 8). To identify potential transcription factor binding sites in the *LIF1* promoter region, we used PlantPAN 3.0 (http://PlantPAN.itps.ncku.edu.tw) (Chow et al., 2015; Chow et al., 2018) “TF/TFBS Search” and “Promoter Analysis”. We also predicted the *SlWRKY16* binding site based on the homologous genes in the phylogenetic tree of WRKY domains on the Plant Transcription Factor Database (PlantTFDB v5.0, http://planttfdb.gao-lab.org<u>; Phylogenetic Tree for Solanum lycopersicum WRKY Family:</u> http://planttfdb.gao-lab.org/phylo_tree.php?sp=Sly&fam=WRKY) (Jin et al., 2016). To determine the consequences of the K251Q amino acid replacement in the LIF1 protein, we predicted potential protein domains of LIF1 using InterProScan (https://www.ebi.ac.uk/interpro/search/sequence/) (Jones et al., 2014; Mitchell et al., 2018). We conducted protein 3D structure prediction and analysis using Phyre2 (Protein Homology/analogY Recognition Engine v 2.0, http://www.sbg.bio.ic.ac.uk/phyre2/html/page.cgi?id=index) (Kelley et al., 2015). We also predicted whether K251Q amino acid substitution is deleterious or neutral using PROVEAN (Choi, 2012; Choi et al., 2012) (Protein Variation Effect Analyzer, http://provean.jcvi.org/seq_submit.php).

### Timecourse RNA-Seq library construction and analysis

We challenged H1706 tomato cultivars with strands of *C. campestris* and collected stem tissues at 1, 2, 3, 4 days post attachment (DPA) and 0 DPA as negative controls. Following this, we constructed strand-specific poly-A based libraries for RNA-seq (Townsley et al., 2015). We conducted sequencing of these libraries on two lanes on Illumina HiSeq 2000 at 50bp Single Read (SR).

We used using CLC Genomics Workbench 11 (QIAGEN) for following RNA-seq analysis. First, we mapped resistant and susceptible cultivar RNA-seq read data to sequenced H1706 tomato genome itag 3.0. To see the general pattern across libraries, we conducted principal component analysis (PCA) of gene expression across different DPA. Next, we used ANOVA comparison with DPA factors and cutoff FDR < 0.1 to select differentially expressed genes (DEG) list. Then, we drew Venn diagrams of DEGs at different DPA libraries. 0 DPA libraries are without *Cuscuta* treatments and serve as the negative control for comparisons. The cutoff of these DEGs are FDR < 0.1 and fold change > 1.5. Following, we constructed a heat map of DEGs across different DPA libraries. Euclidean distance and complete linkage are used for this clustering analysis (Supplemental Figure 1).

### Resistant and susceptible cultivar RNA-Seq library construction and interaction model-based analysis

We challenged the resistant and susceptible tomato cultivars with strands of *C. campestris* and collected stem tissues at 4 days post attachment (DPA). Following this, we constructed strand-specific poly-A based libraries for RNA-seq from the four tomato cultivars, including the resistant cultivars H9492 and H9553, and the susceptible cultivars H9775 and H1706. We conducted sequencing of these libraries on two lanes on Illumina HiSeq 4000 at 100bp Single Read (SR). We first mapped reads to sequenced H1706 tomato genome itag 2.4. To investigate gene expression changes across the resistant and susceptible cultivars in dodder infested versus uninfested plants, we conducted a PCA with K-means clustering using the normalized read counts of sequences mapped to the tomato transcriptome (Supplemental Figure 2). Next, we defined differentially expressed genes with the Bioconductor package DESeq2 employing an interaction design (design = ∼ Condition + Genotype + Condition: Genotype). Following this, we focused on these 113 genes that display expression changes upon dodder infestation that are different in H9492 and H9553 compared to H1706 and H9775. Within these 113 genes, we picked our three candidate genes based on gene annotation and functions.

### Barnes-Hut clustering analysis and gene coexpression network analysis for resistant and susceptible cultivar RNA-Seq

In order to get a more comprehensive differentially expressed gene (DEG) list, we mapped resistant and susceptible cultivar RNA-seq read data to sequenced H1706 tomato genome itag 3.0 by using CLC Genomics Workbench 11 (QIAGEN). Next, we used ANOVA comparison with both factors, all cultivars and with/without *Cuscuta* treatments, and cutoff FDR < 0.1 to select DEG list. In these 10939 genes, we applied Barnes-Hut t-distributed stochastic neighbor embedding (BH-SNE) using RSMod package (script included in code availability) generated 85 gene clusters based on their gene expression patterns. Based on their GO (Gene Ontology) enrichment terms and their included candidate genes, five clusters were selected for further analysis (Supplemental Data Set 9, yellow-labeled genes). We use these selected genes to build gene coexpression networks by using the R script (script included in code availability) that was modified from our previously published method (Ichihashi et al., 2014). We constructed gene coexpression networks for different *C. campestris* treatments in susceptible and resistant cultivars with normal quantile cutoff = 0.997.

### Virus-based Gene Expression (VGE) and virus-induced gene silencing (VIGS) in tomatoes

For preparing the binary vector for plant transient expression that carries the ToMoV DNA, we used a modified pSP72-TAV (Gilbertson et al., 1993; Hou and Gilbertson, 1996) that is lacking the capsid protein ORF (CP) and has a restriction enzyme multisite in which a Gateway® cassette (Thermo Fischer Scientific) was cloned by In-Fusion (Takara) in the *NcoI* site. The whole replicon fragment of TAV-GW was amplified from this vector and cloned into a binary vector to generate a vector for Agrobacterium-mediated transient expression in plants (pMR315). Since the ToMoV DNA-B (Carrying the viral movement protein (MP) and the CP are missing, this clone is not an infectious clone and is only serves as a viral replicon by replicating via rolling circle mechanism (Stenger et al., 1991). The gene cloned into this vector is driven by the CP promoter, which is in the non-translated region between the end of the common region and the start codon of the CP gene that was removed (Supplemental Figure 4).

For transiently overexpressing our candidate genes in the susceptible tomato cultivar H1706, we used this Virus-based Gene Expression (VGE) vector pTAV (Supplemental Figure 4). We cloned *GUS*, *LIF1*, *SlMYB55* and *CuRLR1* genes into pTAV and transformed these into thermo-competent *Agrobacterium tumefaciens* by heat-shock-transformation. For culturing Agrobacterium and preparing agroinjection, we followed the previously published protocol (Vel et al., 2009) with some modifications. For each experiment, we started from growing transformed Agrobacterium on Lysogeny broth (LB) agar plates with appropriate antibiotic selections at 30° C for 2 days. Following this, we inoculated 10 mL liquid LB with transformed Agrobacteria (AGL1) and incubated at 30° C for 16 hours with 200 r.p.m. shaking. We diluted the primary cultures 1:5 into Induction Media (Vel et al., 2009) supplemented with appropriate antibiotic selections and 200 µM acetosyringone, and then incubated them at 30° C for 24 hours with 200 r.p.m. shaking. When the O.D. 600 of the culture was around 1, we harvested the transformed Agrobacteria by centrifuging at 3000 x g for 10 minutes and then resuspended Agrobacteria in Inoculation Buffer (10 mM 2-[N-Morpholino] ethane sulfonic acid (MES), 10 mM MgCl_2_, 200 µM acetosyringone and 0.5 mM dithiothreitol) to an O.D. 600 of 1 culture. Next, we injected this transformed Agrobacterium culture into the first internode of tomato stems using a syringe equipped with a 0.8 mm x 38.1 mm MonoJect needle.

For virus-induced gene silencing (VIGS), we followed the published VIGS in tomato protocol and tobacco rattle virus (TRV)-based vector system (Liu et al., 2002) with slight modifications. This TRV-based VIGS contains TRV-RNA1 (pTRV1) and TRV-RNA2 (pTRV2), which includes multiple cloning sites for building constructs for the genes of interest. Transformed pTRV1 and pTRV2 *Agrobacterium* cultures were mixed in a 1: 1 ratio before infiltration. After about 3.5 hours of incubation on the shaker at room temperature, mixed *Agrobacterium* cultures were infiltrated onto the cotyledons of 9-day-old tomato plants using a 1 ml needleless syringe.

### Preparation of *C. campestris* extracts and injection protocols

For one mL *C. campestris* extracts, we collected 100 mg of the stem tissue in microcentrifuge tubes from *C. campestris* growing on H1706 tomato plants. We used the BioSpec Mini-Beadbeater to grind the liquid nitrogen-frozen tissue with five 2.3 mm diameter BioSpec zirconia beads and 1.0 mm diameter BioSpec zirconia beads in the tubes for 1 minute, and then mixed with one mL deionized water. To remove the plant tissue debris, we centrifuged extracts for 30 seconds at 5000 r.c.f., and used only the supernatant for untreated extract injections. For heat-treated extracts, we heated at 95 °C for 5 minutes. For pH treated extracts, we adjusted the pH to 9 by adding 0.1M NaOH. For filtered extracts, we filtered untreated extracts through a VWR 0.2 µm sterile syringe filter. We injected different treated extracts into the first internode of tomato stems using a syringe equipped with a 0.8 mm x 38.1 mm MonoJect needle. Furthermore, we used 3K, 10K, 30K, and 100K Amicon® Ultra Centrifugal Filter Devices to filter *Cuscuta* extracts. Then, we use flow through extracts to do injection on H9553 stems to test the size of *Cuscuta* signals.

### Protein interaction, subcellular localization and co-localization of fusion candidate proteins

For protein interaction assays, we performed *in vivo* using bimolecular fluorescence complementation (BiFC) system (Kerppola, 2008). The plasmids were constructed by using the Gateway-compatible BiFC vectors SPDK1794 (*p35S::cCitrine*) and SPDK1823 (*p35S::nCitrine*). The leaves of four-week-old *Nicotiana (N.) benthamiana* were injected with different combinations of *Agrobacterium (A.) tumefaciens* GV3101 containing the transient expression vectors (Fang and Spector, 2010).

For subcellular localization and co-localization of fusion candidate proteins, we performed with transient expression fluorescent fusion proteins *in vivo*. The plasmids were constructed by using the Gateway-compatible vectors pGWB5 (*p35S::GFP*) and pGWB660 (*p35S::TagRFP*). The leaves of four-week-old *N. benthamiana* were injected with the *A. tumefaciens* GV3101 strain containing one of the plasmids (Sparkes et al., 2006). To verify the interaction between LIF1 and SlWRKY16 proteins and determine their subcellular localizations when they interact with each other, we also co-expressed the fusion proteins LIF1-GFP (*p35S::LIF1-GFP*) and SlWRKY16-RFP (*p35S::SlWRKY16-TagRFP*). To reduce gene silencing and enhance transient expression L of our candidate proteins, we co-expressed the fusion proteins with p19 obtained from Professor Bo Liu’s Lab at University of California, Davis. Three individual plants and three adult leaves of each plant were used for each treatment. Fluorescence was observed 2 – 5 days after transfection by a Confocal Laser Scanning Platform Zeiss LSM710 (Zeiss, Germany).

### Validation of protein-protein interaction by yeast two-hybrid analysis

To validate the predicted protein-protein interaction between two of our candidate TFs, AP2 and WRKY16, we used the GAL4 yeast two-hybrid system (Clontech). AP2 and WRKY16 were cloned into pGADT7-GW (Addgene Plasmid #61702) and pGBKT7-GW (Addgene Plasmid #61703) plasmids, which were obtained from Yuhai Cui (Lu et al., 2010). Empty pGADT7 (with the GAL4 activation domain, AD) and pGBKT7 (with the GAL4 DNA-binding domain, BD) plasmids were used as negative controls. We use the yeast (*Saccharomyces cerevisiae*) strain AH109 in the MATCHMAKER GAL4 Two-Hybrid System (Clontech), in which *HIS3*, *ADE2*, *MEL1* and *LacZ* are under the control of GAL4 TF. The AD and BD plasmids were co-transformed to yeast AH109 competent cells following Clontech’s user manual instructions of the polyethylene glycol (PGE)/lithium acetate method and cultured in YPD Plus Medium for 90 minutes to promote transformation efficiency. Transformed yeast cells were assayed by culturing in SD/-Leu/-Trp medium to select for successful co-transformants and then assayed by culturing in SD/–Ade/–His/–Leu/–Trp medium with 40 µg/ml X-α-Gal, which provides high-stringency selection. The positive protein-protein interactions between two TFs are indicated by growth on SD/–Ade/–His/–Leu/–Trp/X-α-Gal medium plates and blue colony color.

### Lignin content and composition analysis

Tomato stem material from specified treatments was flash-frozen in liquid nitrogen and lyophilized at -50°C and ≤ 0.1 mbar for 48 hours in a 6L FreeZone 6 Benchtop Freeze Dry System (Labconco Corp., Kansas City, MO, USA). Lyophilized material was pulverized in a 2ml screw top microcentrifuge tube with a glass bead for 10 minutes at a frequency of 20/s in a TissueLyser (Retsch ballmill, Qiagen, Venlo, Netherlands). The material was AIR prepped and destarched using protocols from Barnes and Anderson (Barnes and Anderson, 2017). Acetyl bromide analysis was carried out using the same protocol with one exception: samples were incubated at 50°C with gentle swirling every 10 minutes for 2 hours to limit the degradation of xylan, which can lead to the over quantification of lignin present in the sample. Lignin quantification reactions were performed in a 10 mm light path quartz cuvette (VWR, Radnor, PA, USA, catalog number 414004-062) and absorbance measurements were taken on a SPECTRAmax plus 384 (Molecular Devices, San Jose, CA, USA). Acetyl Bromide Soluble Lignin (%ABSL), which indicates percent absorbance of soluble lignin, was calculated using the extinction coefficient of 17.2 (Chen and Dixon, 2007). Presently, no extinction coefficient is available for tomato stem lignin so the extinction coefficient for tobacco stem lignin was used.

Cell wall-bound aromatics and HPLC analysis of liberated aromatics was performed as described by Eudes et al. (Eudes et al., 2015). The remaining cell wall material post-aromatic extraction, now enriched for lignin, was then washed with water 3 times and dried at 30°C overnight. The chemical composition of the lignin enriched cell wall material was analyzed by pyrolysis-gas chromatography (GC) /mass spectrometry (MS) using a previously described method with some modifications (Eudes et al., 2015). Pyrolysis of biomass was performed with a Pyroprobe 6200 (CDS Analytical Inc., Oxford, PA, USA) connected with GC/MS (GCMS-QP2010 Ultra Gas Chromatograph Mass Spectrometer, Shimadzu corp., Kyoto, Japan) equipped with an AgilentHP-5MS column (30 m 9 0.25 mm i.d., 0.25 lm film thickness). The pyrolysis was carried out at 550 °C. The chromatograph was programmed from 50 °C (1 min) to 300 °C at a rate of 30 °C/min; the final temperature was held for 10 min. Helium was used as the carrier gas at a constant flow rate of 1 mL/min. The mass spectrometer was operated in scan mode and the ion source was maintained at 300 °C. The compounds were identified by comparing their mass spectra with those of the NIST library and those previously reported (Ralph and Hatfield, 1991; Gutiérrez et al., 2006). Peak molar areas were calculated for the lignin degradation products, and the summed areas were normalized per sample.

## Supporting information

Supplemental Data Set 1

Supplemental Data Set 2

Supplemental Data Set 3

Supplemental Data Set 4

Supplemental Data Set 5

Supplemental Data Set 6

Supplemental Data Set 7

Supplemental Data Set 8

Supplemental Data Set 9

## Data availability

All data is available in the main text or the supplementary materials. All DNA-Seq and RNA-Seq raw data are deposited on NCBI SRA PRJNA550259.

## Code availability

All R scripts and package for analysis are deposited on GitHub. R script for RNA-Seq interaction model-based analysis deposited on GitHub (Link: https://github.com/MinYaoJhu/Moran-s-RNA-Seq-analysis-script). R script and RSMod package for Barnes-Hut clustering analysis deposited on GitHub (Link: https://github.com/sdrowland/RSMod). R script for RNA-Seq gene coexpression network analysis deposited on GitHub (Link: https://github.com/Hokuto-GH/gene-coexpression-network-script).

## Author Contributions

M.F. initiated the project with supervision from N. S. and established the connection with Heinz. M.-Y.J. conducted phylogenetic analysis and verified *Cuscuta* species. M.-Y.J., M.F., and R.N.P. conducted histochemical and biomass analysis. M.R. modified the VGE vector and M.F. cloned candidate genes into VGE vectors. M.-Y.J., R.N.P., and L.W. conducted VGE overexpression experiments and analysis for all candidate genes. S.D.R. and M.-Y.J. developed lignin section area quantification methods. M.-Y.J. produced VGE overexpressed samples and M.S.B. performed acetyl bromide, HPLC and Pryo-GC-MS analysis with P.M.S.’s supervision. M.F. made DNA-Seq and RNA-Seq libraries. M.F. conducted preliminary SNP analyses and model-based DEG analyses and picked primary candidate genes. M.-Y.J. performed SNP analyses, protein domain and structure prediction, PROVEAN analysis for DNA-Seq data, and ANOVA DEG analyses, BH-SNE clustering to select candidate genes and built GCNs for RNA-Seq data. S.D.R. developed the RSMod R package. H.N., S.D.R., and M.-Y.J. modified the GCN analysis script and M.-Y.J. construct GCNs. M.-Y.J. and S.D.R. performed PhyloNetwork analysis. M.-Y.J. and L.W. conducted *wrky16* plant phenotyping and VGE overexpression experiments on *wrky16*. M.-Y.J. and L.W. performed subcellular localization and BIFC experiments. M.-Y.J. performed yeast two-hybrid and protein co-localization experiments. M.-Y.J., R.N.P., and L.W. conducted *Cuscuta* extract injection experiments. M.-Y.J. wrote the initial manuscript draft for the main text and made figures and tables with primary editing from N. S.. M.-Y.J., R.N.P., L.W., M.S.B., M. R., and H.N. wrote the initial draft for materials and methods. M.-Y.J., M.F., R.N.P., L.W., M.S.B., H.N., K.Z., P.M.S., and N.R.S. contributed to the editing of this manuscript. N.R.S. supervised the whole project.

## Acknowledgments

We are grateful to S. Dinesh-Kumar, A. B. Britt, S. Brady, and D. Runcie for their input on this research, and S. Dinesh-Kumar, A. B. Britt, S. Brady, and E. D. Marable for editing suggestions. We thank C. Wong, C. Ito, Z. Jaramillo, J. Tanurahardja and J. Lu for helping with some parts of experiments or image analysis, and Axtell Lab and N. Johnson for providing information to help with verifying *Cuscuta* species, and Liu Lab and Y. Lee for providing p19, pGWB5 and pGWB660 plasmid. We also thank R. Ozminkowski, Manager of Agricultural Research from HeinzSeed for providing seeds of three Heinz cultivars, and the UC Davis Plant Transformation Facility for generating transgenic plants, and CyVerse for online data storage. This work used the Extreme Science and Engineering Discovery Environment (XSEDE) on the Pittsburgh Supercomputing Center (PSC) at the Regular Memory (Bridges) servers through allocation IBN200015. We thank A. Eudes for his recommendations and guiding conversations in lignin analysis, and J. Ortega for her technical support with the pyro-GC/MS instrument.

