## Supplemental Figure 1 to 21 for "Lignin-based resistance to *Cuscuta campestris* parasitism in Heinz resistant tomato cultivars"

**RESEARCH ARTICLE**

**This PDF file includes:**

Supplemental Figure 1 to 21

Legends for Supplemental Data Set 1 to 9

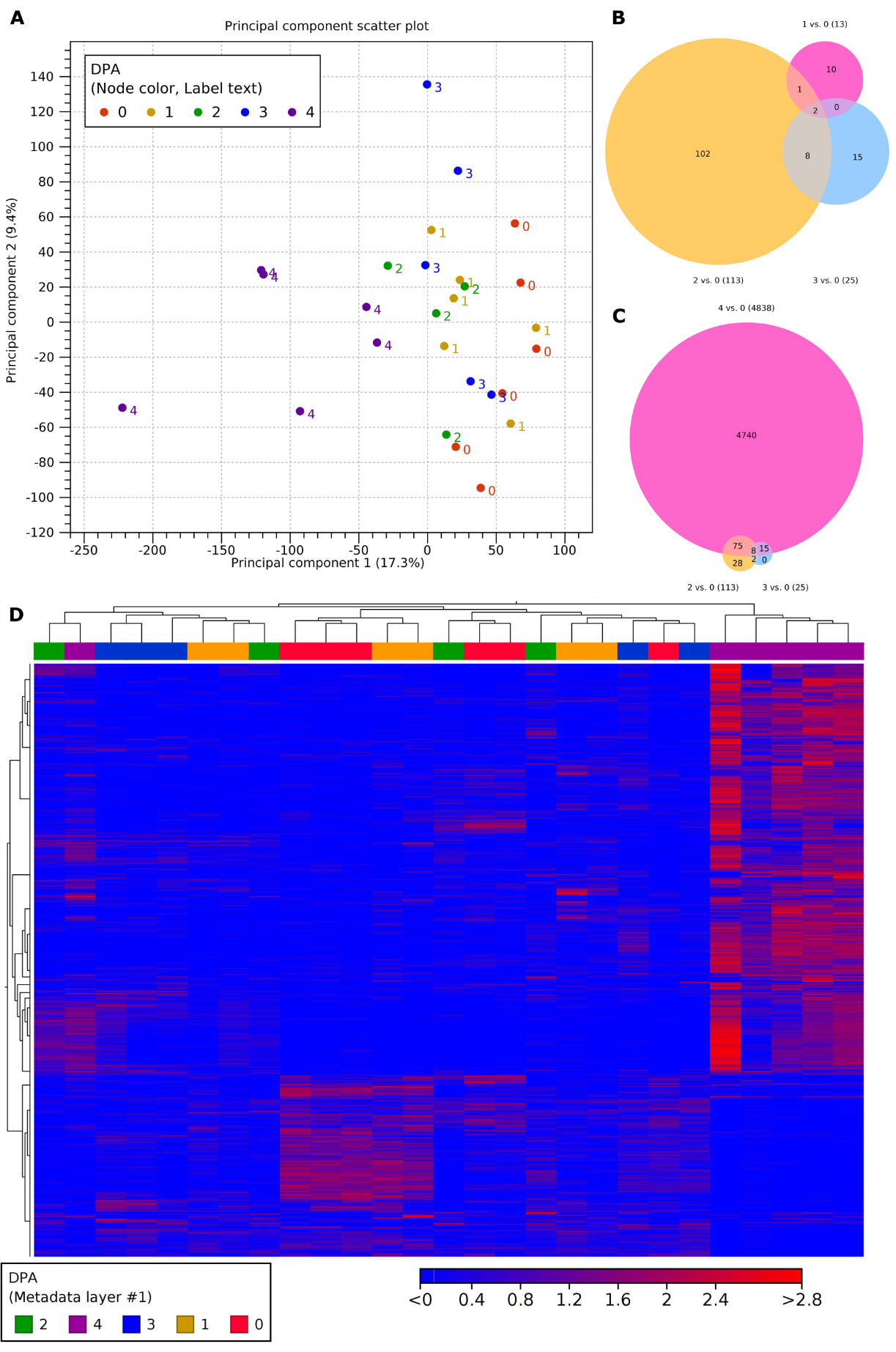

Supplemental Figure 1 | Analysis of time-course RNA-Seq data. (A) Principal component analysis (PCA) of gene expression across different day post attachment (DPA). Library number, 0 PDA, n = 6; 1 DPA, n = 6; 2 DPA, n = 4; 3 DPA, n = 5; 4 DPA, n = 6. (B-C) Venn diagram of differentially expressed genes (DEGs) at different DPA libraries. 0 DPA libraries are without *Cuscuta* treatments and serve as the control for comparisons. The cutoff of these DEGs are FDR < 0.1 and fold change > 1.5. (D) Heat map of DEGs across different DPA libraries. DEGs are selected by ANOVA analysis with cutoff FDR < 0.1. Euclidean distance and complete linkage are used for this clustering analysis.

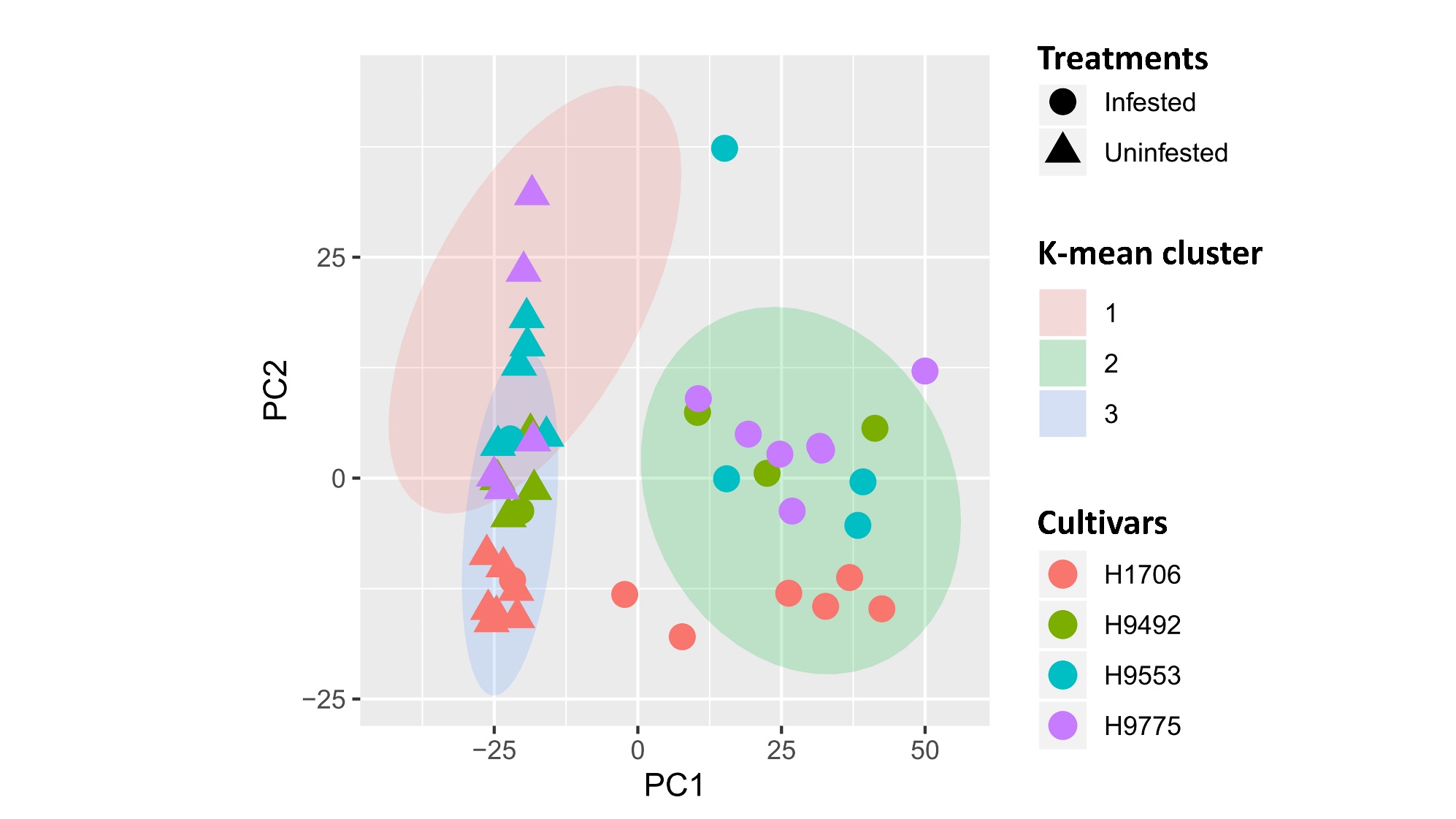

Supplemental Figure 2 | RNA-Seq analysis results of gene expression across resistant and susceptible cultivars at 4 DPA. PCA of gene expression across resistant and susceptible cultivars at 4 DPA. Different treatment conditions are represented by shapes: square dots indicate the uninfested host stem tissue samples; circle dots indicate the infested host stem tissue samples. Different cultivars are represented by different colors.

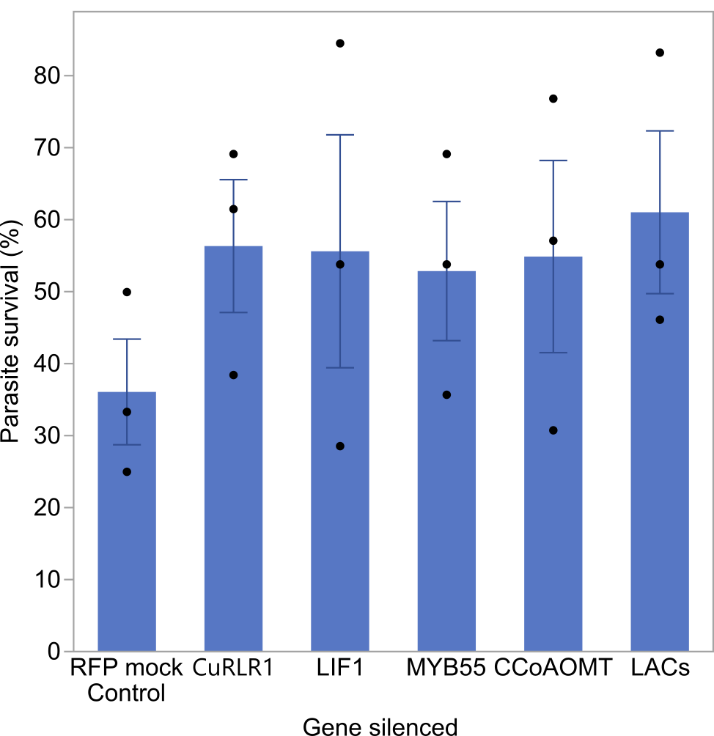

Supplemental Figure 3 | Virus-induced gene silencing (VIGS) in resistant tomato H9553. Plot shows the average *C. campestris* survival rates on different VIGS tomato plants. Survival rates = (the number of *C. campestris* surviving by the end of the experiment/ the total number of *C. campestris* at the beginning of experiment) * 100%. Each dot represents the average survival rate for each experiment, which used 12-14 individual plants (biological replicates) for each VIGSed gene. CCoAOMT, caffeoyl-CoA O-methyltransferase; LAC, laccase.

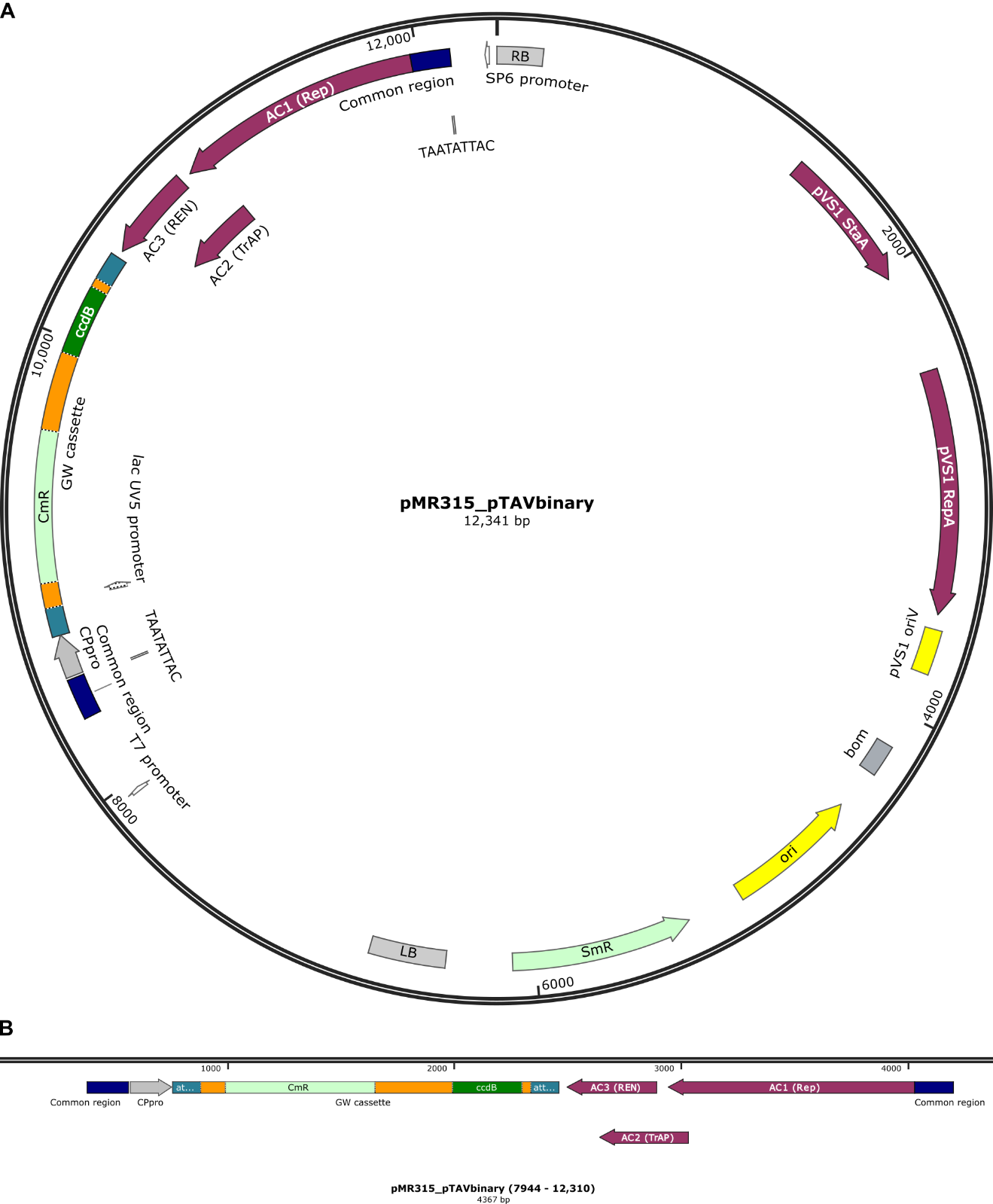

Supplemental Figure 4 | Virus-based Gene Expression vectors (pTAV) map. (A) Full map of pTAV plasmid. (B) Zoom-in view of the Gateway® cassette region. The gene cloned into this vector is driven by the capsid protein promoter (CPpro), which is in the non-translated region between the end of the common region and the start codon of the capsid protein gene that was removed. CmR indicates chloramphenicol acetyltransferase, which is the Chloramphenicol resistance gene. SmR indicates aminoglycoside adenylyltransferase, which is the Spectinomycin/Streptomycin resistance gene. pVS1 RepA indicates replication protein from the *Pseudomonas* plasmid pVS1. pVS1 StaA indicates stability protein from the *Pseudomonas* plasmid pVS1. Complete sequence of pTAV is attached in Datasets S1.

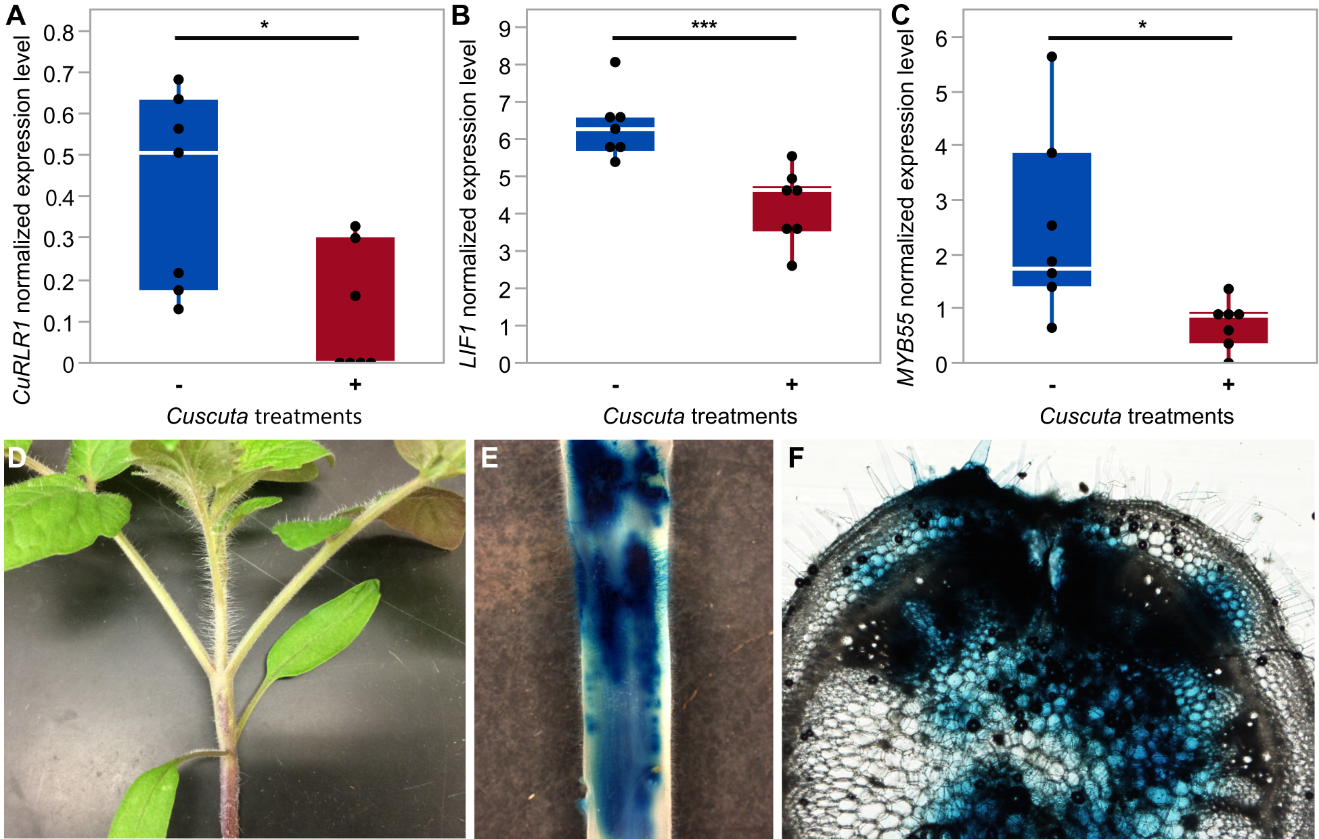

Supplemental Figure 5 | Expression of candidate genes and Virus-based Gene Expression (VGE) of GUS in tomato H1706. (A-C) The normalized expressions levels (CPM, counts per million) of genes in susceptible cultivar H1706 under *C. campestris* infestation. – and + indicates without or with *C. campestris* infection treatments respectively. Biologically independent replicates: RNA-Seq libraries: H1706-Cc, n = 7; H1706+Cc, n = 7. Data are assessed using two-tailed t test. “*”: p-values < 0.05, “**”: p-values < 0.01, “***”: p-values < 0.005. (D) Tomato seedling showing first internode. Arrow points to injection site. (E) The first internode of stem stained for GUS expression from VGE construct in susceptible cultivar H1706. (F) Hand section (about 300 μm) of stem near injection site stained for GUS expression.

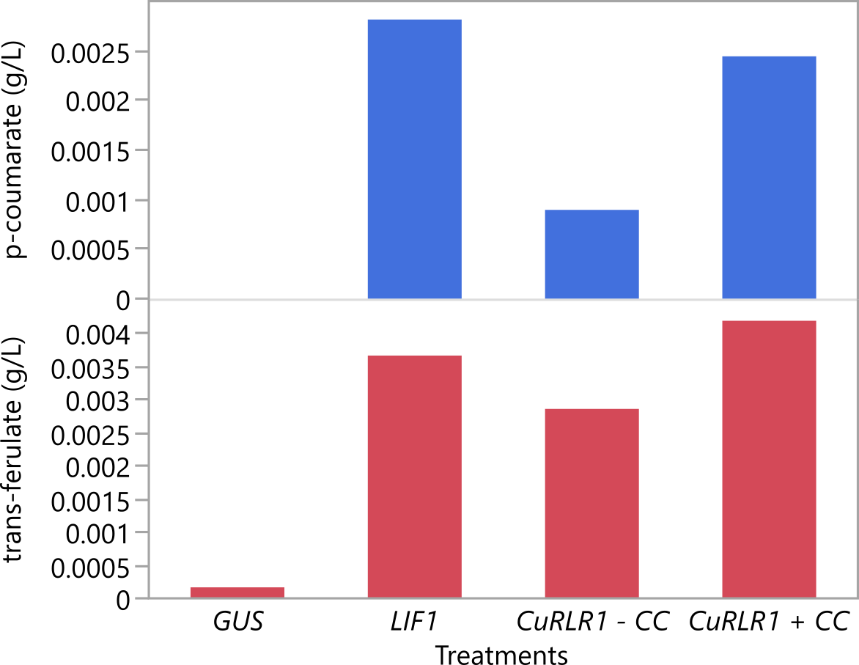

Supplemental Figure 6 | HPLC data for p-coumarate and trans-ferulate levels in different VGE and *C. campestris* infection treatments. HPLC data for p-coumarate and trans-ferulate was generated from ethyl acetate extract of de-starched AIR prepped stem tissue. The unit of this data is g/L. –Cc and +Cc indicate without or with *C. campestris* infection treatments respectively. Biological replicates collected from first internodes; *GUS*, n = 8; *LIF1*, n = 8, *CuRLR1*-Cc, n = 18; *CuRLR1*+Cc, n = 18. HPLC assay technical replicate, n = 1.

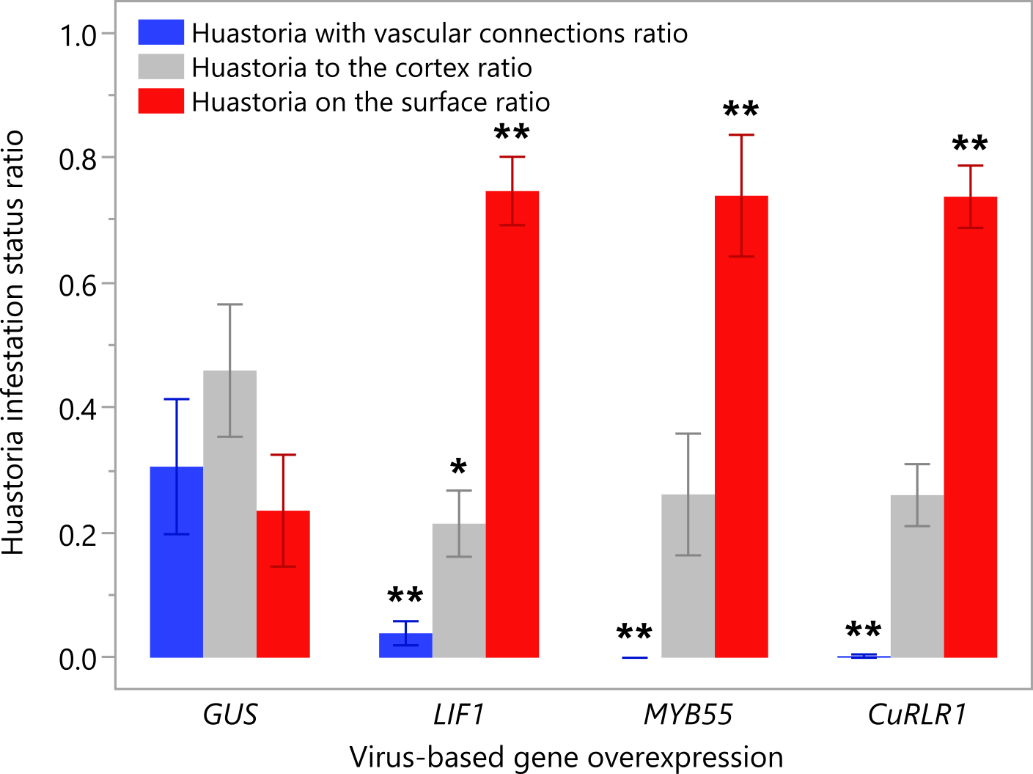

Supplemental Figure 7 | Haustorium infestation status ratio under different VGE treatments. The numbers of haustoria in different infestation status are quantified by examining hand sections and vibratome sections. The ratios are calculated by dividing the number of haustoria in each status by the total number of haustoria on each section. The detailed haustorium number and ratio data are presented in Supplemental Data Set 3. Data are analyzed using Dunnett's test. “*”: p-values < 0.05, “**”: p-values < 0.01.

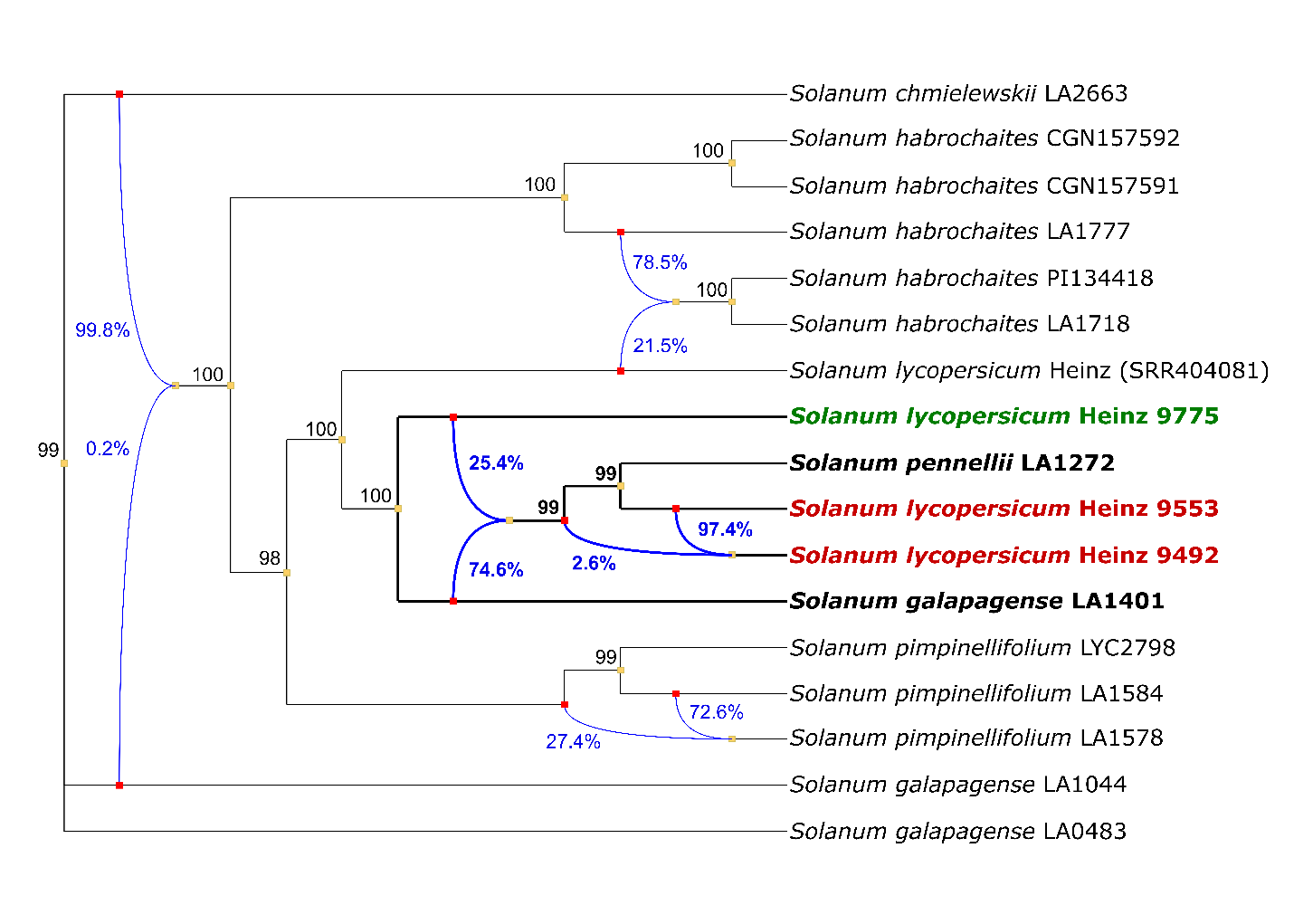

Supplemental Figure 8 | Phylogenetic network analysis using 500 kb sequence around the *LIF1* resistance-specific SNP enriched region. This phylogenetic network analysis was conducted using PhyloNetworks in the Julia environment with 500 kb of sequence around the *LIF1* resistance-specific SNP enriched region (SL3.0 ch02: 43800000 – 44300000). The species and accessions relevant to this work are highlighted in bold front and darker lines. Blue lines indicate potential hybridization events among these tomato cultivars, accessions, and species. Blue percentage numbers represent the gene flow from each potential parent cultivar to the hybridization event. Black numbers next to each node are bootstrap values. Green bold labeled cultivar H9775 is susceptible to *C. campestris* infection. Red bold labeled cultivars H9553 and H9492 are resistant to *C. campestris* infection. Black bold labeled species are potential tomato wild species introgression sources contributing to the *LIF1* resistance-specific SNP enriched region.

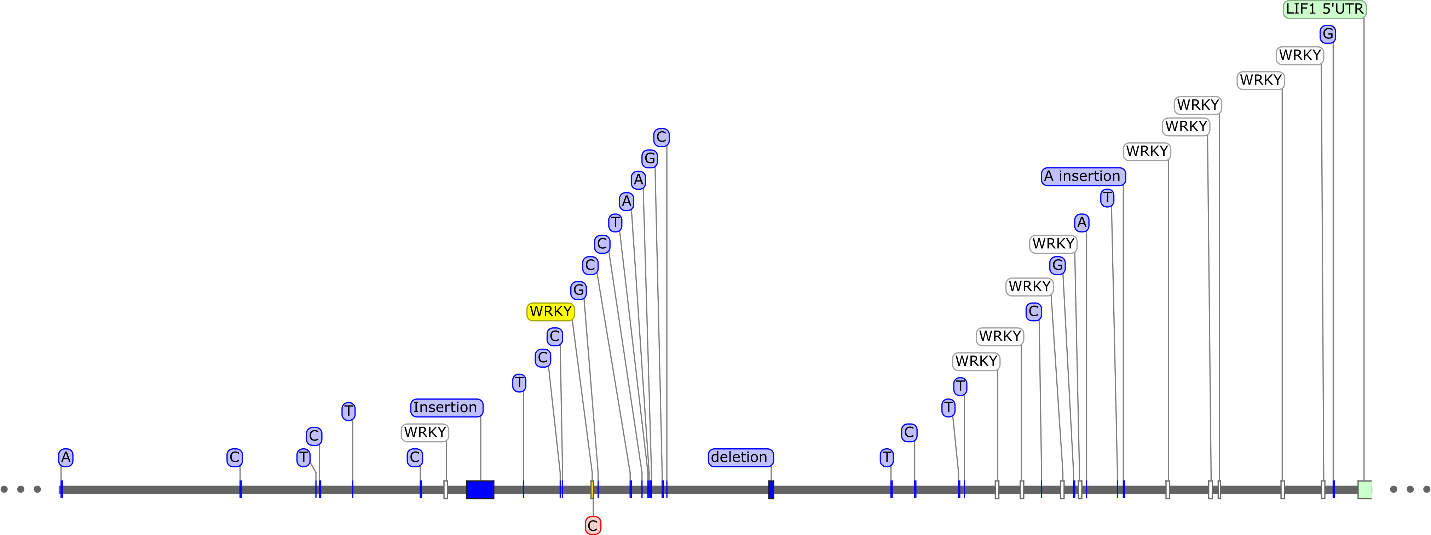

Supplemental Figure 9 | *LIF1* promoter region and transcription factors binding motifs. The *LIF1* promoter DNA sequence is labeled as a gray colored line. Transcription factors binding motifs are labeled as white boxes on the DNA sequence, and the potential key WRKY binding site is labeled as a yellow box. Resistance specific SNPs are labeled as blue lines on the DNA sequence. The resistance specific SNP that is located on the key WRKY binding site is labeled in red color.

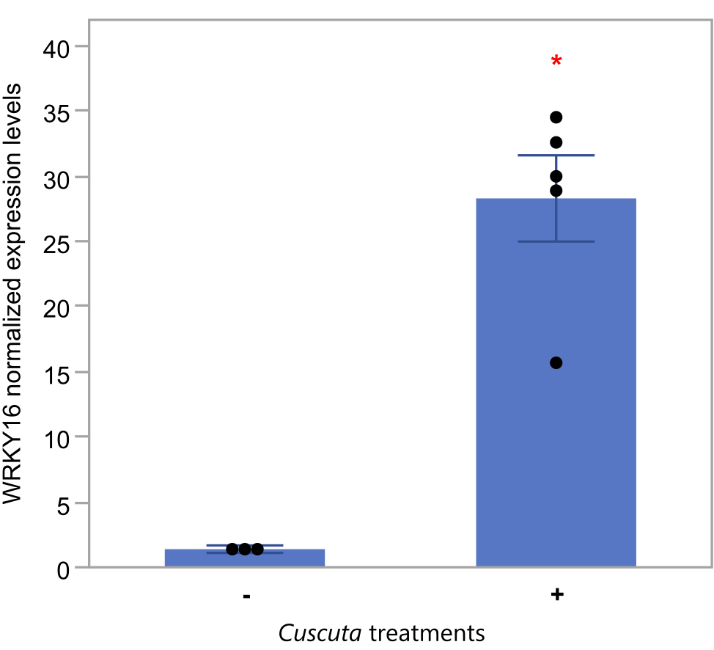

Supplemental Figure 10 | *SlWRKY16* expression level in different *Cuscuta* treatment condition. Normalized *SlWRKY16* expression level from qPCR data in M82 tomatoes with/without *Cuscuta* treatments. – and + indicates without or with *C. campestris* infection treatments respectively. Biologically independent replicates: M82-Cc, n = 3; M82+Cc, n = 5. Data presented are assessed using two-tailed t test. “*”: p-values < 0.01. Value of the t-statistic: 8.10; degrees of freedom: 4.06.

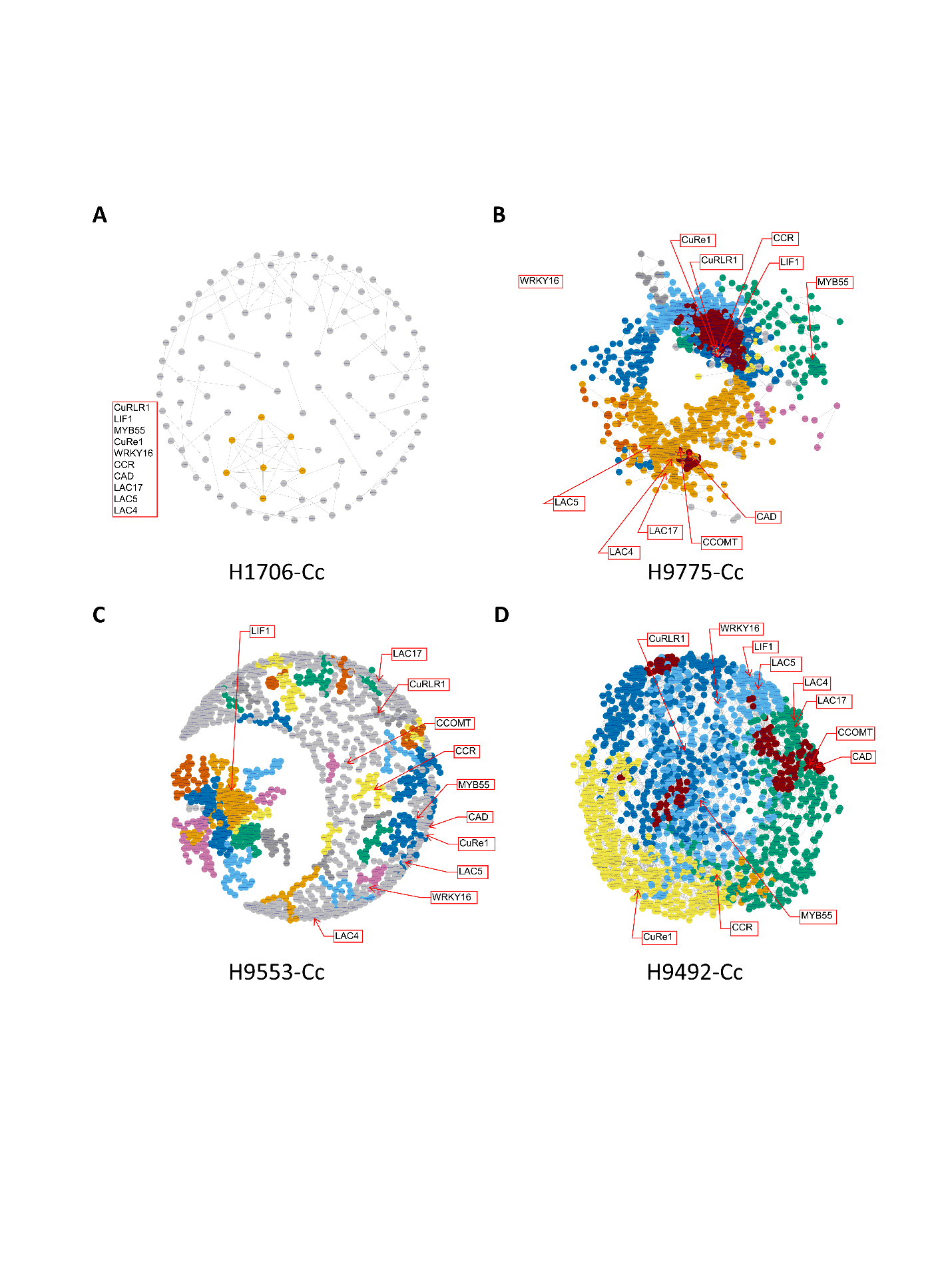

Supplemental Figure 11 | Gene co-expression network (GCN) analysis of identified key regulators. Gene co-expression networks (GCNs) of four different Heinz susceptible and resistant cultivars without *C. campestris* treatments. Based on BH-SNE analysis, 1676 genes in cluster 11, 17, 23, 39, 46 and CuRLR1 are selected for building GCNs. -Cc indicate without *C. campestris* infection treatments. The genes that are listed at the left of the GCN and not labeled in the network are the genes that have no coexpression connection with all the other genes in list.

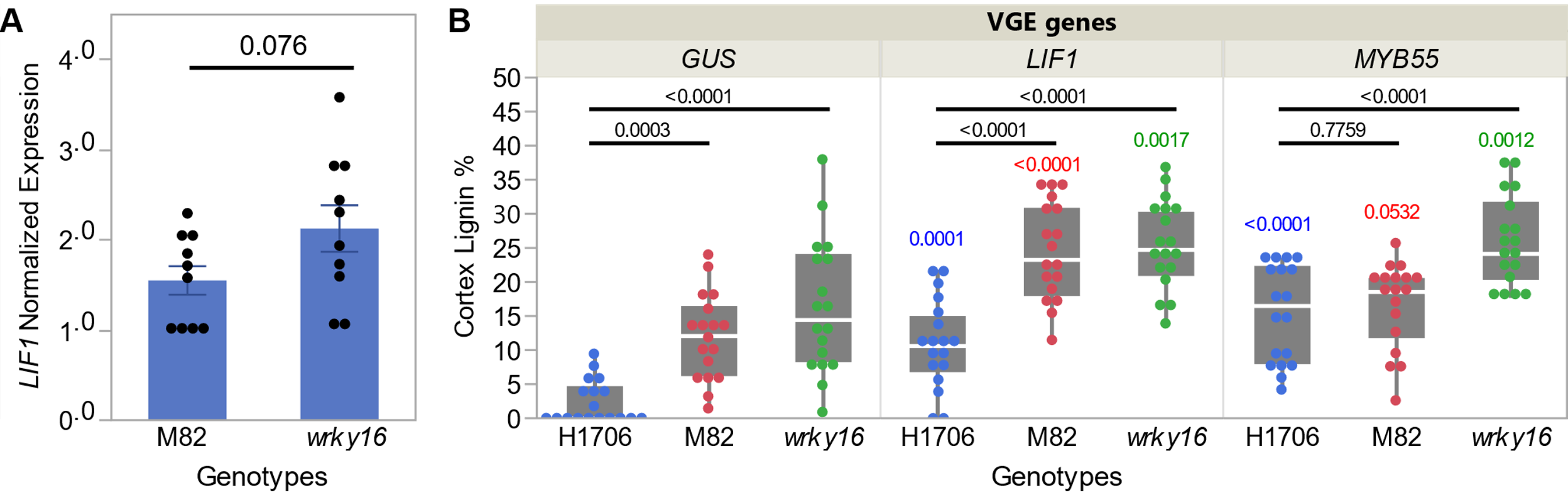

Supplemental Figure 12 | *LIF1* expression levels in *wrky16* and VGE overexpressing *SlMYB55* and *LIF1* in H1706, M82, and *wrky16*. (A) Normalized *LIF1* expression level from qPCR data in M82 and *wrky16* tomatoes. Data presented are assessed using Student's t test and p-value is labeled above the boxplot. Replicates: n = 10 for each plant genotype. (B) VGE overexpressing *SlMYB55* and *LIF1* in both susceptible H1706 and M82 tomatoes, and resistant *wrky16*. Data presented are assessed using Dunnett's test with H1706 as the control in each VGE overexpressing group and p-values are labeled above the boxplot in black. Data presented are also assessed using Dunnett's test with *GUS* as the negative control for each plant genotype. Each plant genotype group is labeled in specific color (H1706 in blue, M82 in red, *wrky16* in green) and p-values are labeled above the boxplot in the corresponding color. Replicates: n = 18 for each treatment.

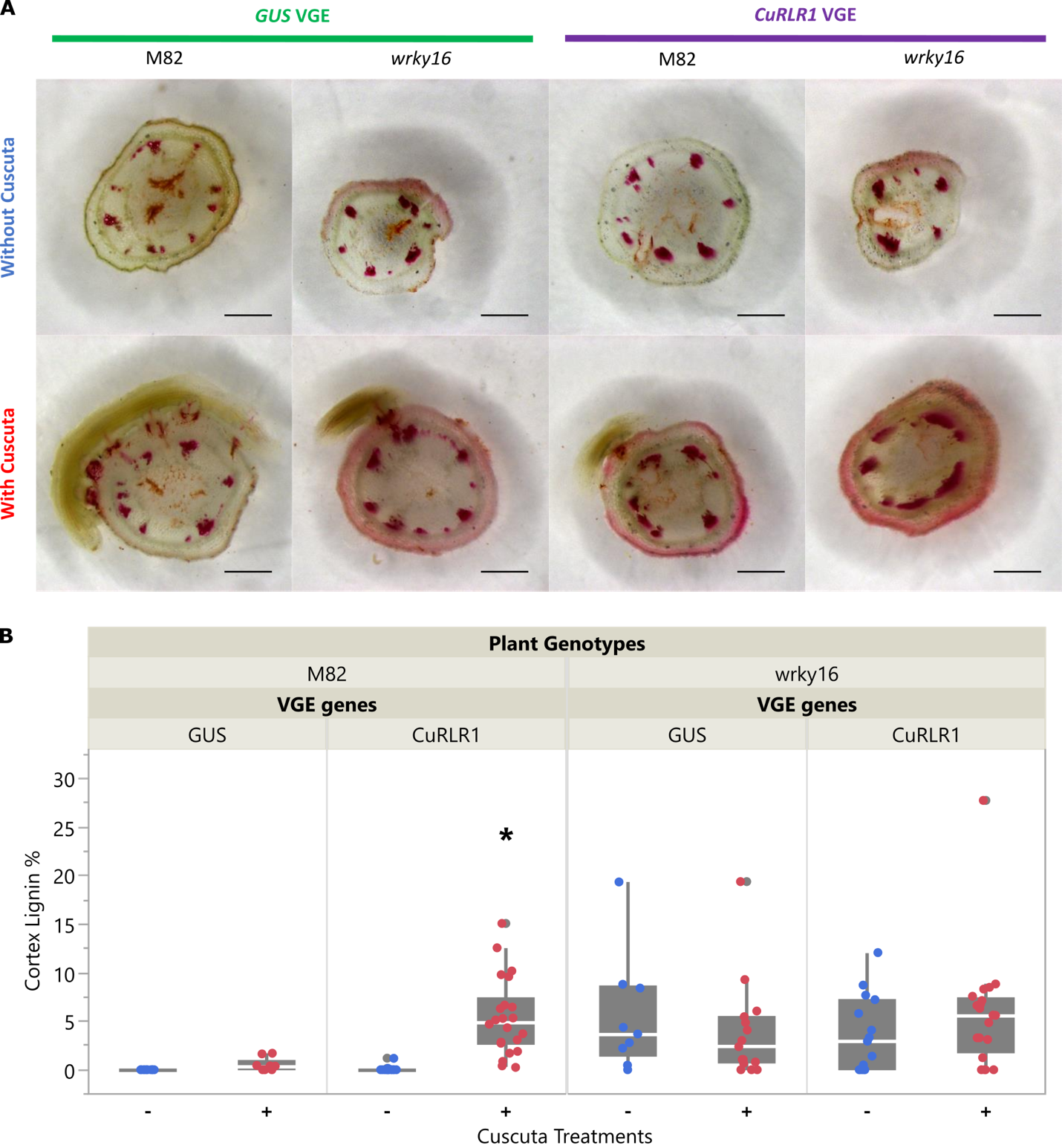

Supplemental Figure 13 | VGE overexpressing *CuRLR1* induced stem lignification in susceptible M82 tomatoes but not in *wrky16* tomatoes. (A) ~300 μm sections of the haustoria attachment sites stained with Phloroglucinol-HCl. Scale bar, 1 mm. (B) Cortex lignin area percentage in both susceptible M82 tomatoes and wrky16 tomatoes. – and + indicates without or with *C. campestris* infection treatments respectively. Data are assessed using Dunnett's test with *GUS*-Cc as the negative control for each genotype. “*”: p-values < 0.01. Replicates: M82+*GUS*-Cc, n = 10; M82+*GUS*+Cc, n = 9; M82+*CuRLR1*-Cc, n = 16; M82+*CuRLR1*+Cc, n = 22; *wrky16*+*GUS*-Cc, n = 9; *wrky16*+*GUS*+Cc, n = 15; *wrky16+CuRLR1*-Cc, n = 15; *wrky16+CuRLR1*+Cc, n = 20. Samples were collected at 10 DPI and 7 DPA.

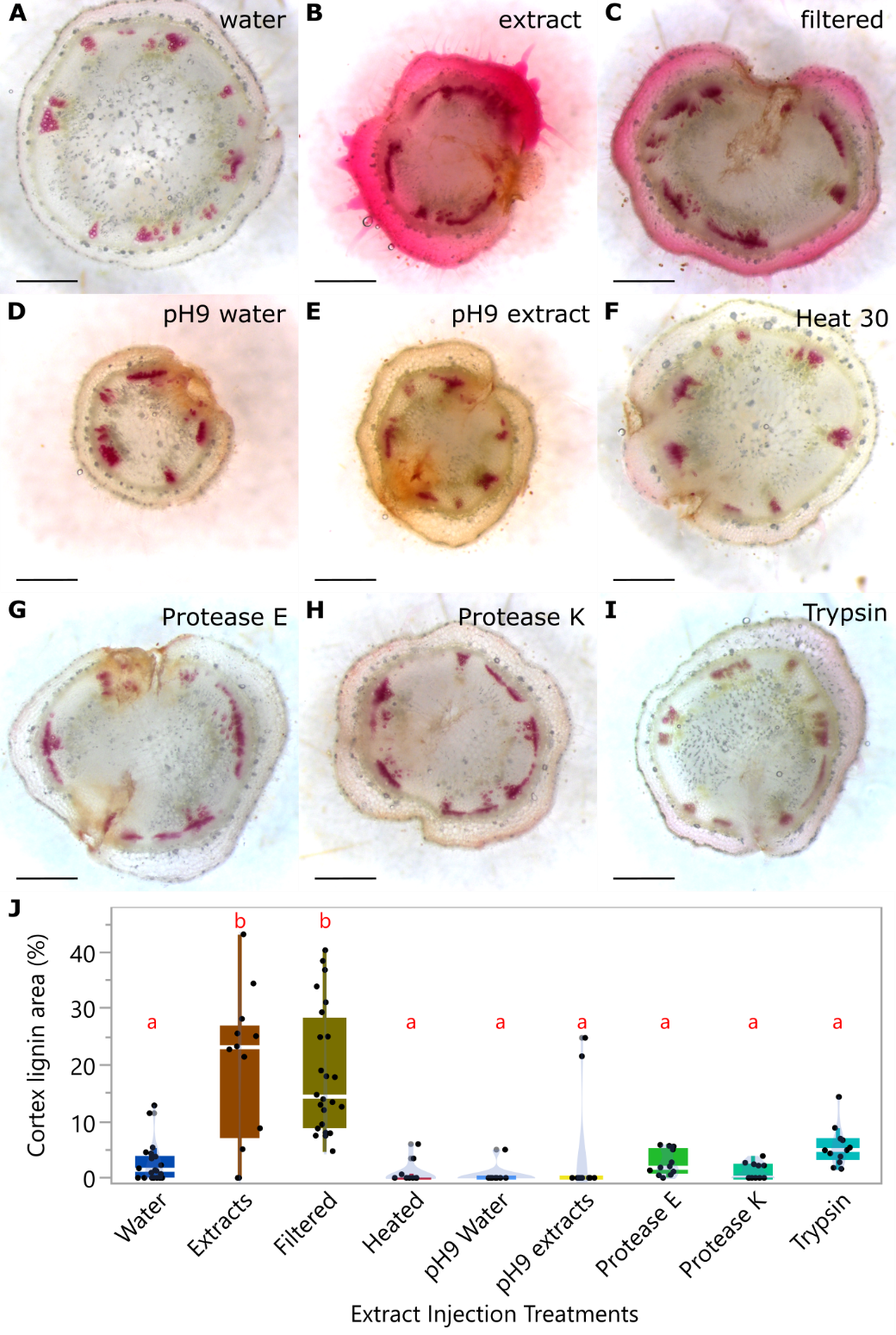

Supplemental Figure 14 | *C. campestris* extract injections to detect *Cuscuta* signals. (A-I) ~300 μm hand sections of resistant H9553 stems near injection sites stained with Phloroglucinol-HCl. Lignin is stained red. Data were collected at 7 days post injection (DPI). The H9553 plants are injected with (A) water, (B) untreated *C. campestris* extract, pH 5.8, (C) *C. campestris* extract filtered with 0.2 µm filter, (D) pH 9 water, and (E) pH 9 *C. campestris* extract, (F) heat-treated *C. campestris* extract (95°C for 30 minutes), (G) Protease E-treated *C. campestris* extract, (H) Protease K-treated *C. campestris* extract, and (I) Trypsin-treated *C. campestris* extract. (J) Percentage of lignified cortex area in total stem area. The samples injected with water serve as negative controls. Different treated or untreated *C. campestris* extracts are compared to negative controls. Data presented are assessed using pair-wise comparisons with Tukey test. P-value of the contrasts between “a” and “b” are less than 0.01. Replicates: water, n = 22; untreated extract, n = 13; filtered extract, n = 24; heat-treated extract, n = 12; pH 9 water, n = 11; pH 9 extract, n = 12; Protease E-treated, n = 12; Protease K-treated, n = 12; Trypsin-treated, n = 12.

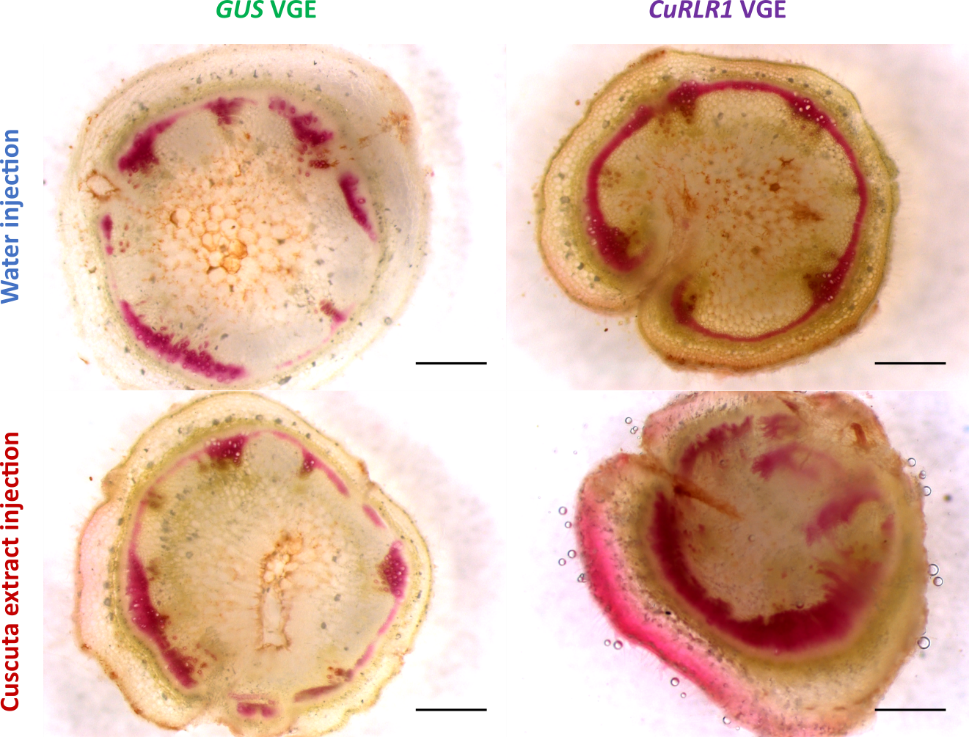

### Supplemental Figure 15 | VGE overexpressing CuRLR1 in H1706 with or without Cuscuta extract injections. These are ~300 μm sections of the haustoria attachment sites stained with Phloroglucinol-HCl. Scale bar, 1 mm. VGE overexpressing GUS served as a negative control for VGE. Water injection functions as a negative control for extract injections.

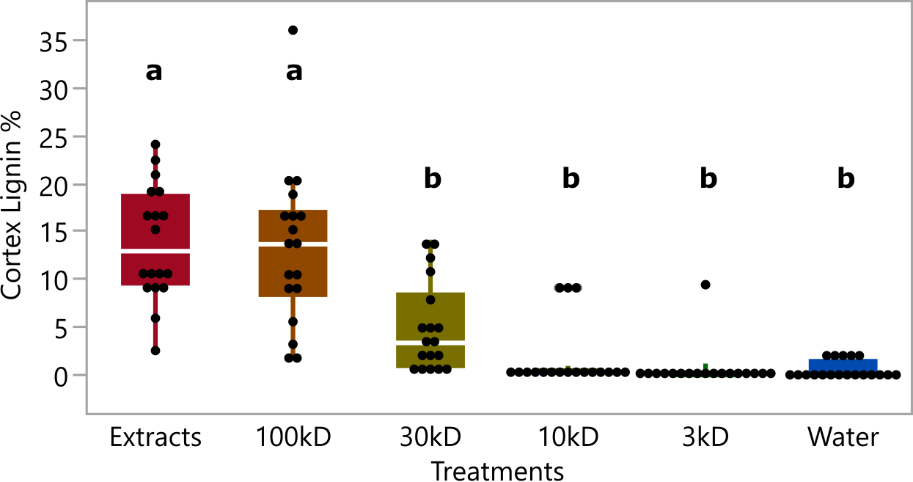

### Supplemental Figure 16 | Cuscuta signal size analysis by using Cuscuta extract injections. Cortex lignin area percentage in H9553 cultivar with different size filtered Cuscuta extract injection. The Cuscuta extracts that flow through 3kD, 10kD, 30kD, and 100kD Amicon® Ultra Centrifugal Filter Devices are used to do injection on H9553 stems to test the size of Cuscuta signals. Data were assessed using pair-wise comparisons with Tukey test. P-values between “a” and “b” are < 0.05. Replicates: untreated extracts, n = 18; 100kD filtered extract, n = 18; 30kD filtered extract, n = 18; 10kD filtered extract, n = 18; 3kD filtered extract, n = 18; water, n = 19.

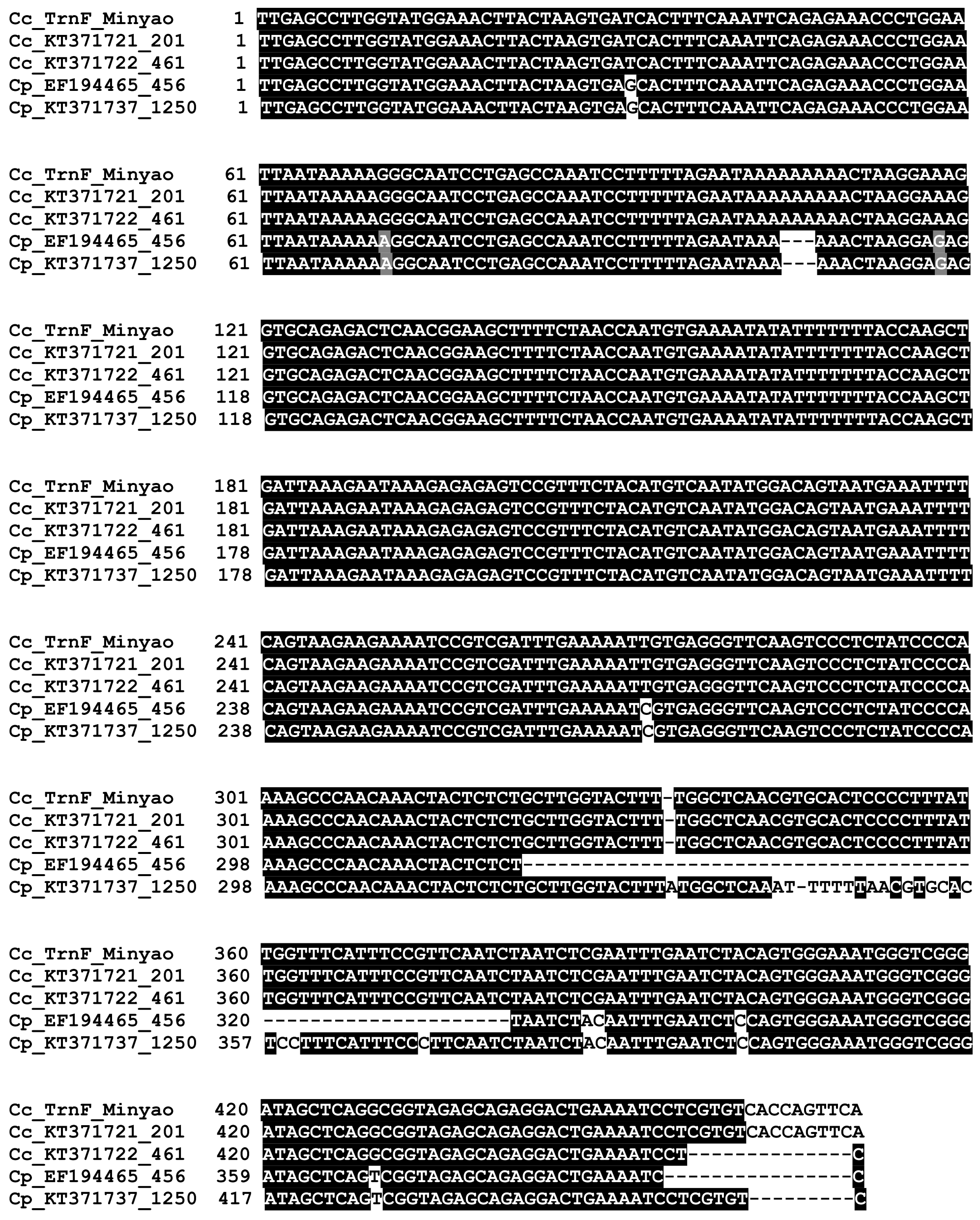

### Supplemental Figure 17 | Sequence alignment of plastid trnL-F intron / spacer region sequences in our Cuscuta campestris isolate and published Cuscuta campestris and Cuscuta pentagona. The first sequence is from the Cuscuta campestris isolate we used in this research. The other sequences are from previously published trnL-F sequences of Cuscuta campestris (Cc) and Cuscuta pentagona (Cp) with GenBank accession numbers and following by DNA accession numbers.

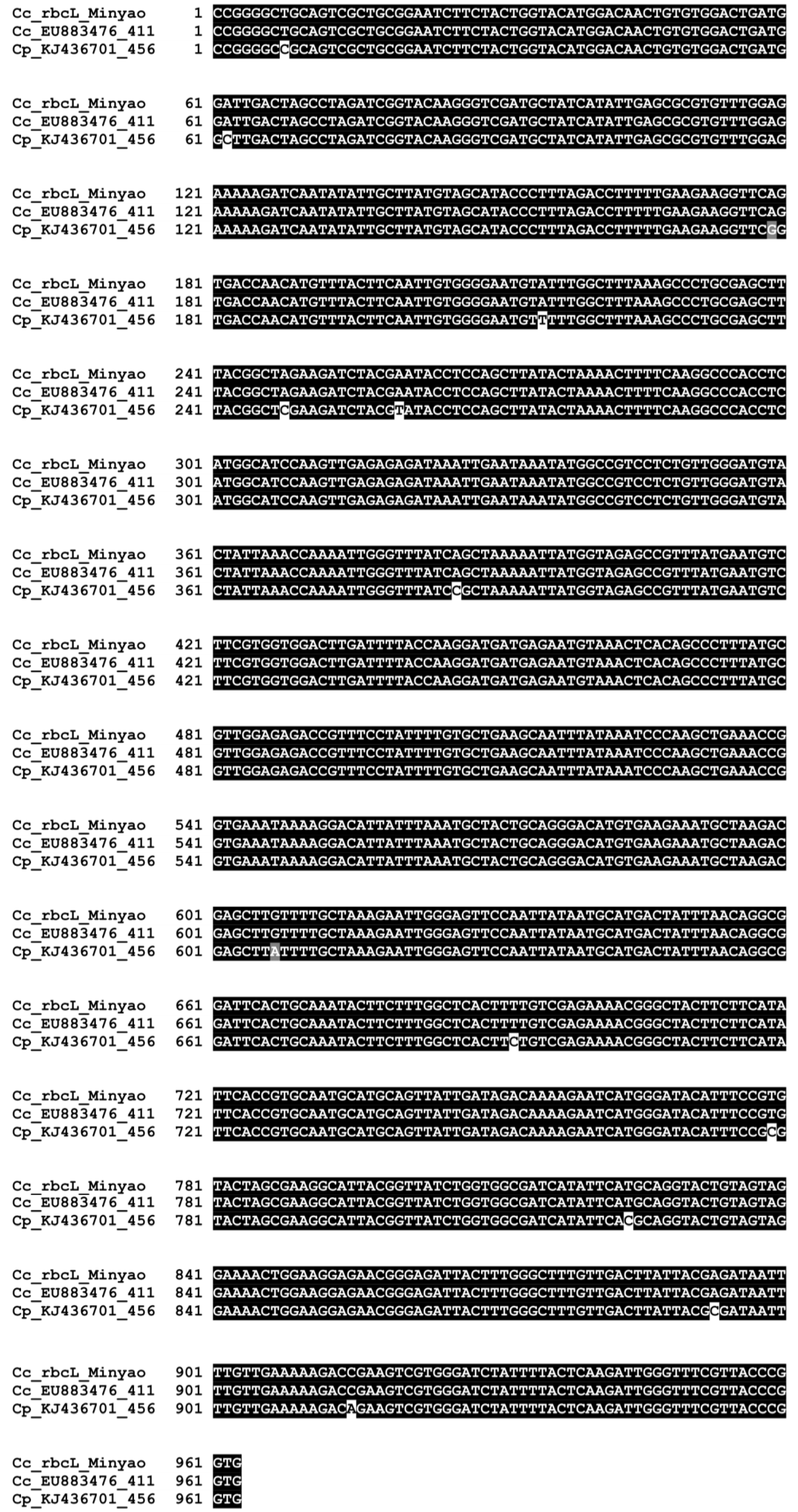

### Supplemental Figure 18 | Sequence alignment of plastid ribulose-1,5-bisphosphate carboxylase/oxygenase large subunit (rbcL) sequences in our Cuscuta campestris isolate and published Cuscuta campestris and Cuscuta pentagona. The first sequence is from the Cuscuta campestris isolate we used in this research. The other sequences are from previously published rbcL sequences of Cuscuta campestris (Cc) and Cuscuta pentagona (Cp) with GenBank accession numbers and following by DNA accession numbers.

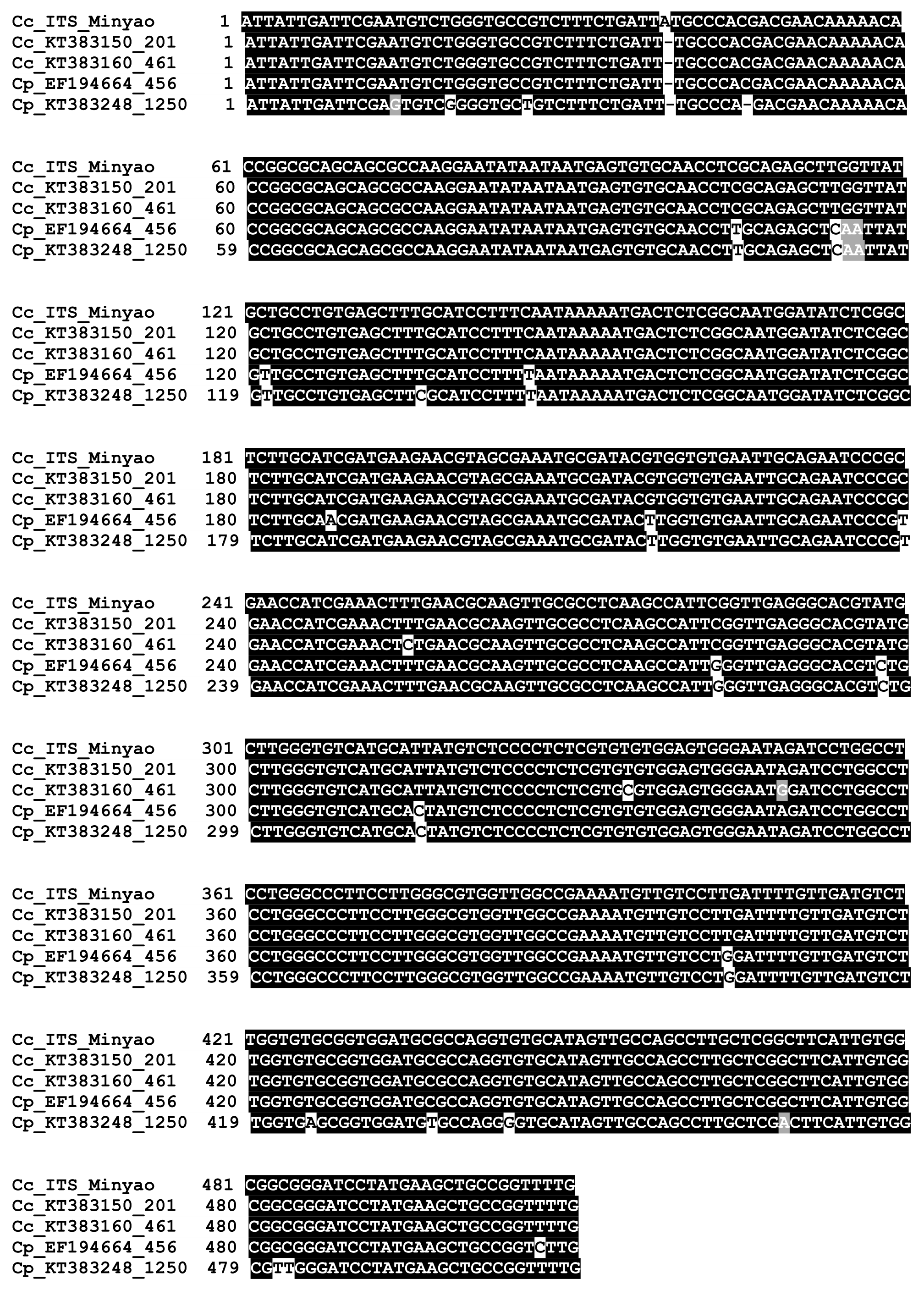

### Supplemental Figure 19 | Sequence alignment of nuclear internal transcribed spacer (nrITS) sequences in our Cuscuta campestris isolate and published Cuscuta campestris and Cuscuta pentagona. The first sequence is from the Cuscuta campestris isolate we used in this research. The other sequences are from previously published nrITS sequences of Cuscuta campestris (Cc) and Cuscuta pentagona (Cp) with GenBank accession numbers and following by DNA accession numbers.

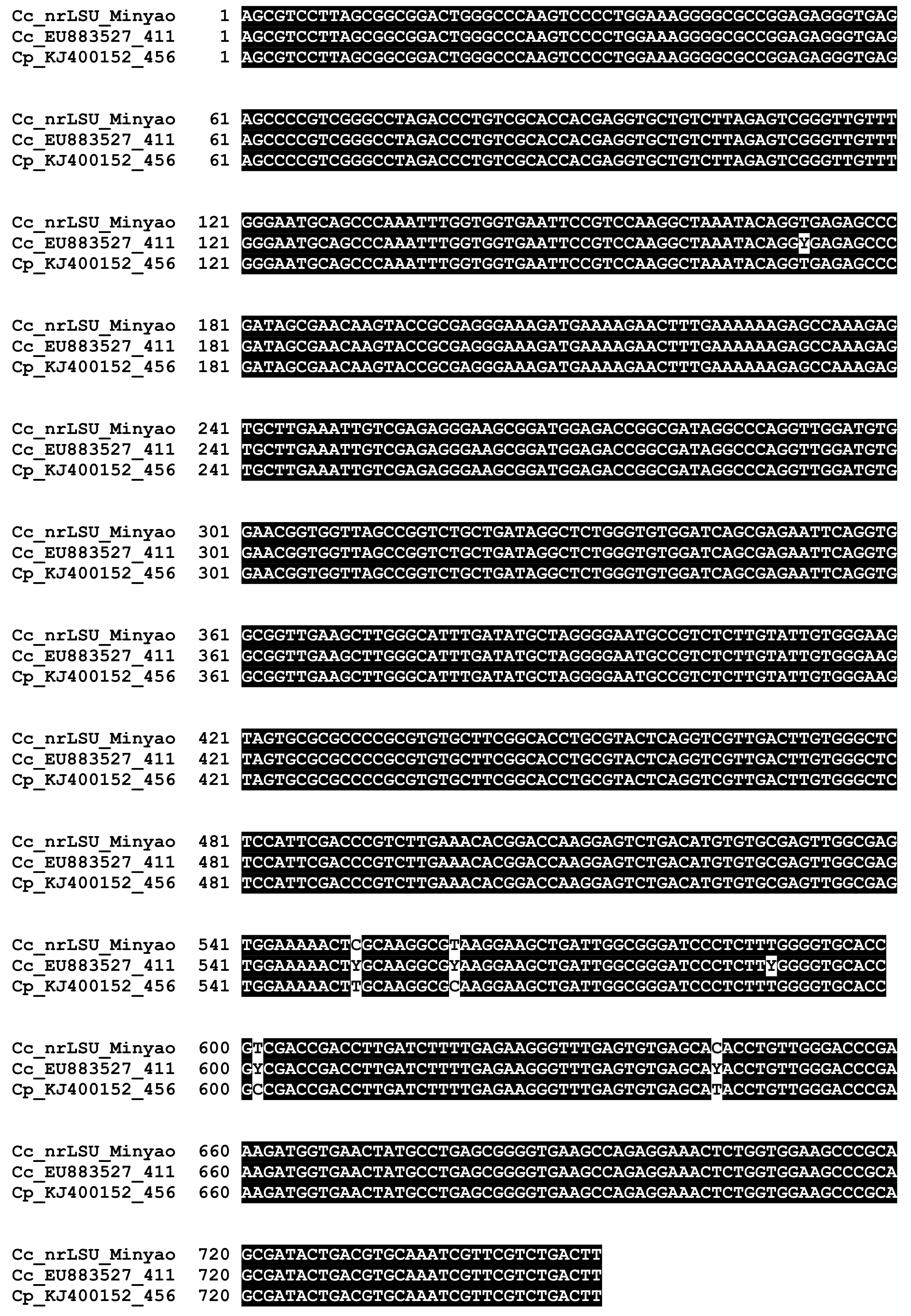

### Supplemental Figure 20 | Sequence alignment of nuclear large-subunit ribosomal DNA (nrLSU) sequences in our Cuscuta campestris isolate and published Cuscuta campestris and Cuscuta pentagona. The first sequence is from the Cuscuta campestris isolate we used in this research. The other sequences are from previously published nrLSU sequences of Cuscuta campestris (Cc) and Cuscuta pentagona (Cp) with GenBank accession numbers and following by DNA accession numbers.

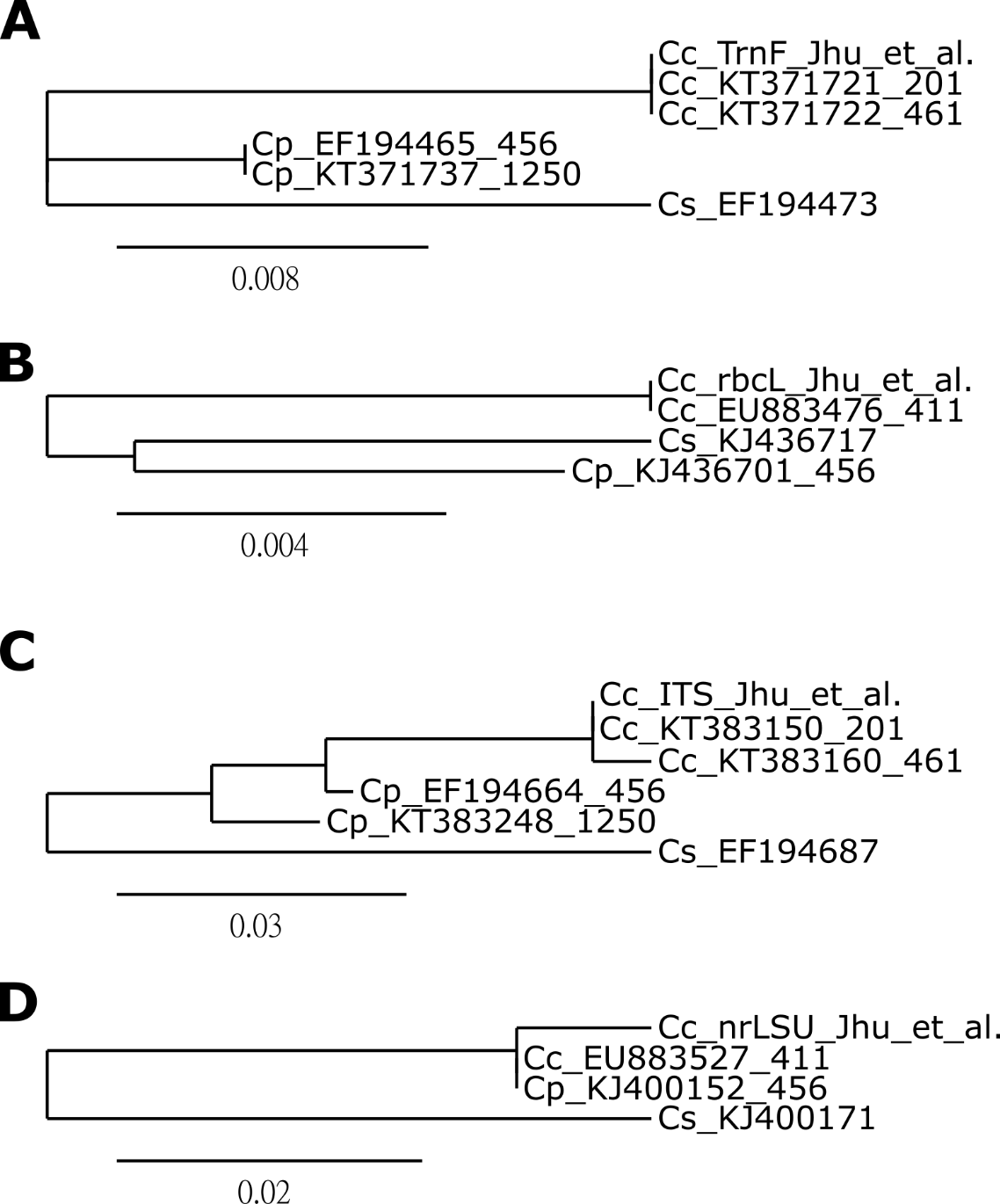

### Supplemental Figure 21 | Phylogenetic relationships among our Cuscuta campestris isolate and published Cuscuta campestris and Cuscuta pentagona by Maximum-Likelihood Phylogenies. The first sequence of each tree is from the Cuscuta campestris isolate we used in this research. The other sequences are from previously published (A) TrnL-F (B) rbcL (C) nrITS (D) nrLSU sequences of Cuscuta campestris (Cc) and Cuscuta pentagona (Cp) with GenBank accession numbers and following by DNA accession numbers. These trees are rooted using C. stenolepis (Cs) as functional outgroup. The number on the scale represents the percentage of genetic variation.

### Supplemental Data Set 1. DEG list of time-course RNA-Seq data.

Supplemental Data Set 1 provided as an Excel file.

### Supplemental Data Set 2. DEG list of resistant and susceptible host response to C. campestris by using an interaction design model.

Supplemental Data Set 2 provided as an Excel file.

### Supplemental Data Set 3. Vector pTAV (pMR315_pTAV-GW binary) sequence

Supplemental Data Set 3 provided as a text file.

### Supplemental Data Set 4. Haustorium infestation status quantification.

Supplemental Data Set 4 provided as an Excel file.

### Supplemental Data Set 5. Resistant specific SNPs in all chromosome.

Supplemental Data Set 5 provided as an Excel file.

### Supplemental Data Set 6. Resistant specific SNPs in LIF1 promoter region.

Supplemental Data Set 6 provided as an Excel file.

### Supplemental Data Set 7. Predicted transcription factor binding sites in LIF1 promoter region.

Supplemental Data Set 7 provided as an Excel file.

### Supplemental Data Set 8. DEG list of four different Heinz tomato cultivars response to C. campestris by ANOVA analysis.

Supplemental Data Set 8 provided as an Excel file.

### Supplemental Data Set 9. Gene list in the Barnes-Hut t-distributed stochastic neighbor embedding (BH-SNE) generated clusters.

Supplemental Data Set 9 provided as an Excel file.
